# OX40 agonism enhances efficacy of PD-L1 checkpoint blockade by shifting the cytotoxic T cell differentiation spectrum

**DOI:** 10.1101/2021.12.24.474145

**Authors:** Guillaume Beyrend, Tetje C. van der Sluis, Esmé T.I. van der Gracht, Tamim Abdelaal, Simon P. Jochems, Robert A. Belderbos, Thomas H. Wesselink, Suzanne van Duikeren, Floortje J. van Haften, Anke Redeker, Elham Beyranvand Nejad, Marcel Camps, Kees LMC Franken, Margot M. Linssen, Peter Hohenstein, Noel F.C.C. de Miranda, Hailiang Mei, Adriaan D. Bins, John B.A.G. Haanen, Joachim G. Aerts, Ferry Ossendorp, Ramon Arens

## Abstract

Immune checkpoint therapy (ICT) has the potency to eradicate cancer but the mechanisms that determine effective *versus* non-effective therapy-induced immune responses are not fully understood. Here, using high-dimensional single-cell profiling we examined whether T cell states in the blood circulation could predict responsiveness to a combined ICT, sequentially targeting OX40 costimulatory and PD-1 inhibitory pathways, which effectively eradicated syngeneic mouse tumors. Unbiased assessment of transcriptomic alterations by single-cell RNA sequencing and profiling of cell-surface protein expression by mass cytometry revealed unique activation states for therapy-responsive CD4^+^ and CD8^+^ T cells. Effective ICT elicited T cells with dynamic expression of distinct NK cell and chemokine receptors, and these cells were systemically present in lymphoid tissues and in the tumor. Moreover, NK cell receptor-expressing CD8^+^ T cells were also present in the peripheral blood of immunotherapy-responsive cancer patients. Targeting of the NK cell and chemokine receptors in tumor-bearing mice showed their functional importance for therapy-induced anti-tumor immunity. These findings provide a better understanding of ICT and highlight the use of dynamic biomarkers on effector CD4^+^ and CD8^+^ T cells to improve cancer immunotherapy.

## INTRODUCTION

Immunotherapy has become an important treatment option for cancer patients, but is only effective in a minority of patients. Therefore, a deeper understanding of factors governing immune responses upon immunotherapy is required to extend clinical efficacy to the majority of patients (1). Many studies have focused on characterizing intratumoral CD8^+^ T cells (2), however system-wide profiling studies have demonstrated that systemic anti-tumor immune responses are essential for immunotherapeutic efficacy (3). A comprehensive description of how effective cancer immunotherapy affects the T cell states in the blood circulation is currently lacking.

Regression of solid tumors is generally positively correlated with T cells infiltrating the tumor tissue (4). Expression of inhibitory molecules including PD-1 and CTLA-4 on these T cells, however, is associated with impaired function such as diminution of the cytotoxic and proliferative potential (5). Moreover, T cell costimulation is often diminished in tumor settings (6), leading to suboptimal T cell activation. To counteract the cancer-associated T cell inhibition, successful immunotherapies named immune checkpoint therapy (ICT), were developed that blocked the inhibitory molecules PD-1 and CTLA-4 (7, 8). Immune checkpoint blockade provides currently a recognized treatment option for several types of cancer (1, 9). However, response rates are still low, immune-related adverse events occur frequently and long-term survival can only be achieved in a minority of patients (10), which warrants to determine the probability of a clinical response and the development of more efficacious treatment options. In this respect, a better understanding of the cellular mechanisms that mediate tumor rejection will support both the clinical prospects as the design of optimal treatment modalities (11). Moreover, predictive biomarkers related to effective therapy are highly desired, especially in light of the numerous clinical trials with novel (combinatorial) immunotherapeutic approaches that are on-going (12-14). In addition, methods directly targeting costimulatory receptors, such as members of the TNF receptor superfamily (e.g. CD27, CD134 (OX40) and CD137 (4-1BB), expressed on tumor-specific T cells have been developed, and showed potential by itself and combined with immune checkpoint blockade (15, 16), but mechanisms underlying the combinatorial ICT are lacking.

Emerging single-cell technologies such as single-cell RNA sequencing (scRNA-seq) and high-dimensional flow and mass cytometry has provided unprecedented insight into the heterogeneity of the tumor microenvironment (TME) and its modulation by immunotherapy (17-21). For example, these single-cell technologies highlighted the identification of intratumoral T cells with different states of functionality ranging from cytotoxic to dysfunctional (22) and the existence of biomarkers in CD8^+^ T cells that associated with responsive tumor regression (23). Predictive biomarkers in patients treated with ICT such PD-1 and CTLA-4 blockade have also been investigated in the systemic circulation, showing key roles for T cells (24-33), NK cells (34), and monocytes (35, 36).

Here, we performed deep profiling of the systemic T cell response induced by immunotherapeutic regimens built on driving agonist signals via OX40-mediated costimulation in conjunction with inhibition of negative costimulation by the PD-1-PD-L1 pathway. The additive effects of combination therapy over monotherapy were assessed by studying the transcriptional and proteomic changes of therapy-responsive T cell populations using two complementary high-dimensional single-cell profiling techniques, i.e. scRNA-seq (20) and mass cytometry (37). We found that the combined ICT elicited most profoundly effector T cell subsets in the blood with dynamic kinetics, and which were characterized by specific biomarkers (e.g. transcription factors, cytotoxic molecules, chemokines and NK cell markers). System-wide analysis revealed that the therapy-responsive T cells were not limited to the blood but connected to other key immune compartments like the spleen, bone marrow and lymph nodes. Analysis of human peripheral blood mononuclear cells obtained shortly after PD-1 therapy, revealed similar effector T cell states in the blood that correlated with the clinical response rate. The identified biomarkers resembled a closely related set of genes, which functionally connected to the treatment efficacy. Together, this study reveals dynamic cellular changes occurring during effective ICT, and the role of a set of biomarkers connected to the cytotoxic potential of CD8^+^ T cells, which are instrumental in tumor immunity and could be used to assess the level of immunotherapy efficiency.

## RESULTS

### Identification of circulating immunotherapy-responsive T cell subsets by single-cell transcriptional profiling

To examine whether stimulating costimulatory receptors can improve PD-(L)1 checkpoint blockade, we challenged wild-type mice with syngeneic MC-38 tumors (38), which represents an ICT sensitive colorectal tumor model, and treated tumor-bearing mice with anti-PD-L1 antibodies, blocking the inhibitory PD-1/PD-L1 pathway, and with agonistic antibodies targeting the costimulatory receptor OX40 **(Figure 1A)**. Blockade of the PD-1/PD-L1 axis resulted in delayed tumor outgrowth, whereas anti-OX40 treatment did not a significantly delay tumor outgrowth **(Figure S1A, S1B)**. Moreover, addition of the TLR9 ligand CpG augmented the antitumoral actions of anti-OX40 **(Figure S1A, S1B)**, which is in line with a previous report (39). Strikingly, the combination of PD-L1 blockade and anti-OX40/CpG treatment cured the majority of mice **(Figure 1B)**. The combination of the two immunotherapeutics (anti-PD-L1 and anti-OX40/CpG), referred hereafter as PDOX, was also most effective against established syngeneic HCmel12 melanoma tumors (40) **(Figure 1B)**.

**Figure 1.**
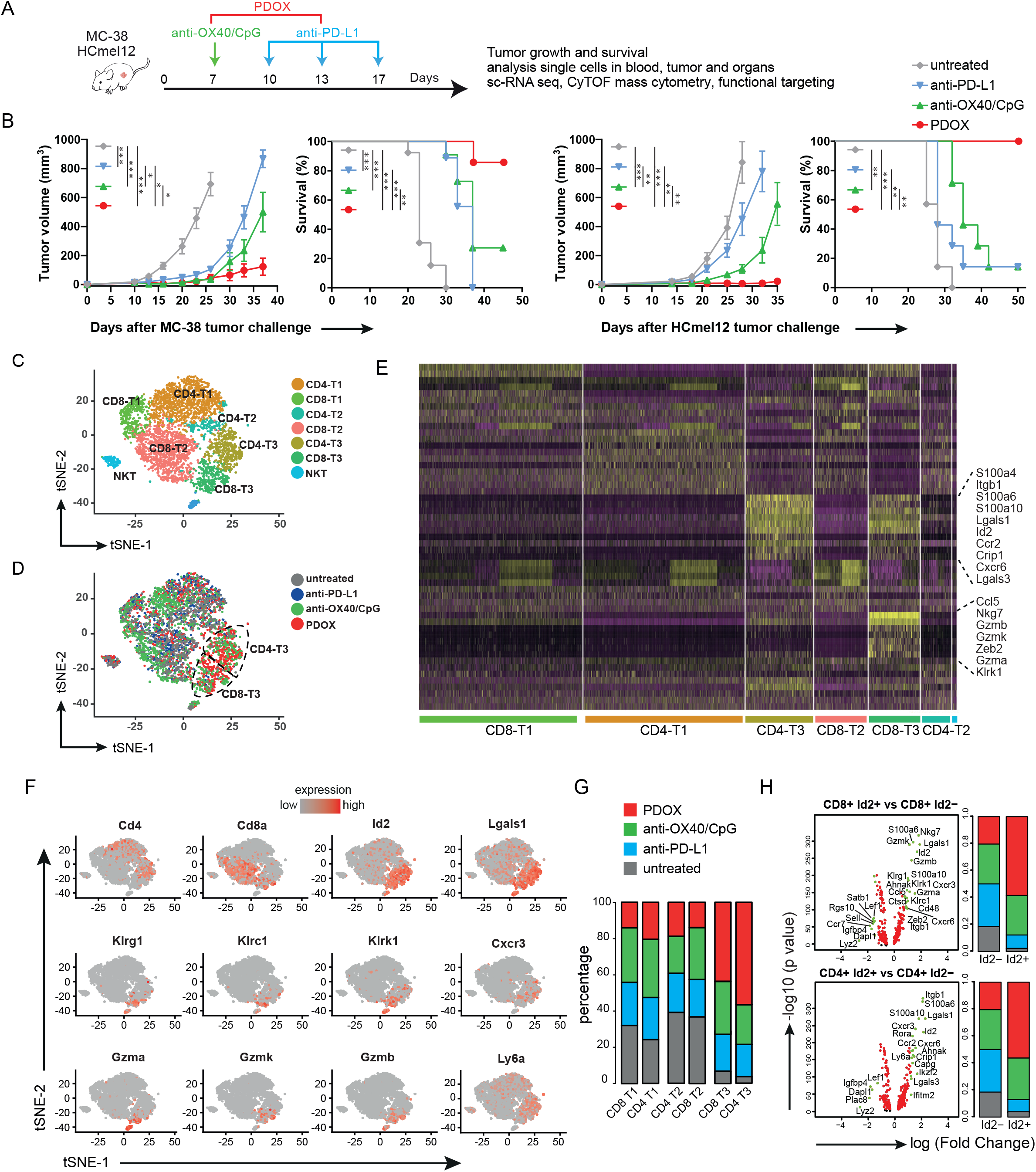
Transcriptional profiling identifies therapy-responsive T cell subsets in the blood circulation. (A) Schematic of the immune checkpoint therapy (ICT) regimen strategy. C57BL/6 mice were subcutaneously challenged with MC38 or HCmel12 syngenic tumors and treated with different ICTs. (B) MC-38 and HCmel12 tumor growth (mean ± SEM) and survival curves of untreated (grey), anti-OX40/CpG (green), anti-PD-L1 (blue), and anti-OX40/CpG plus anti-PD-L1 (termed PDOX) treated wild-type mice. Data were combined from two replicate experiments (n=8-16 mice per group). *p<0.05, **p<0.01, ***p<0.001 (C) t-SNE (t-distributed stochastic neighbor embedding) scRNAseq plot visualizing six transcriptional clusters of blood T cells from day 18 tumor-bearing mice that were untreated or received ICT. (D) t-SNE scRNAseq plot of blood T cells color coded for the untreated and ICT groups. (E) Heatmap displaying scaled expression values of discriminative genes per cluster. (F) t-SNE scRNAseq plots displaying gene expression of *Cd4, Cd8, Id2, Lgals1, Klrg1, Klrc1, Klrk1, Cxcr3, Gzma, Gzmk, Gzmb, Ly6a*. (G) Stacked bar graphs representing the percentage of cells from the untreated and ICT groups present in the six transcriptional clusters. (H) Volcano plots showing significant gene expression related to Id2, Klrk1, and Klrc1 expression in CD4^+^ and CD8^+^ T cells. The log2 fold change (FC) in gene expression on the x-axis and unadjusted P values on the y-axis are illustrated. Black dots represent genes with adjusted P-value > 0.05, red dots represents genes with adjusted P-value < 0.05 and absolute average log2 Fold-Change < 1, green dots with gene name represent genes with adjusted P-value < 0.05 and absolute average log2 Fold-Change> 1. Stacked bar graphs indicate the percentage of the cell origin according to their treatment.

To identify circulating T cell subsets that associate with effective checkpoint therapy, we isolated CD4^+^ and CD8^+^ T cells from the peripheral blood at day 18 post MC-38 tumor challenge of treated and untreated mice. Per condition >1,000 cells were analyzed by single-cell RNA sequencing (scRNA-seq) with a coverage of 60,000 reads per cell. The subpopulation structure of the circulating T cells was defined by pooling data from the different treatment groups representing 5,600 cells total and using the Seurat package analysis to identify transcriptional clusters. Six distinct T cell clusters could be identified consisting of three CD4^+^ and three CD8^+^ T cell clusters **(Figure 1C-1E)**. Two clusters (CD4-T3, CD8-T3) were over-represented in the PDOX group **(Figure 1D-1G)**, and both these T3 clusters were characterized by *Id2* and *Lgals1* transcripts encoding for the transcription factor ID2 and Galectin-1, respectively **(Figure 1E,1F)**. Other gene transcripts overrepresented in both CD4-T3 and CD8-T3 clusters were *Cxcr3* (coding for the chemokine receptor CXCR3) and *Ly6a* (coding for Sca-1) **(Figure 1F)**. Transcripts linked to NK cell markers and cytotoxicity including *Klrk1* (coding for NKG2D protein), *Klrc1* (coding for NKG2A), *Klrg1* (coding for KLRG1), *Gzma, Gzmb*, and *Gzmk* (coding for Granzyme A, B, K) were mostly enriched in CD8-T3 (**Figure 1F**).

Expression of ID2 as well as of KLR familymembers are linked to cytotoxic effector CD8^+^ T cell differentiation but their interconnectivity is unknown (41-43). To gain more insight into this connection of the *Id2*^*+*^, *Klkr1*^*+*^, and *Klrc1*^*+*^ subsets within the T3 clusters, the transcriptional profile of these subsets within the CD8^+^ and CD4^+^ T cell lineages were analyzed distinctly **(Figure 1H, Figure S1C)**. Upregulation of *Id2* transcripts in CD8^+^ T cells associated with the transcripts of *Klrk1, Klrc1, Klrg1, Nkg7, Gzma, Gzmb, Gzmk, Zeb2, S100a6, S100a10, Lgals1, Cxcr3, Cxcr6, Cd48, Ctsd, Itgb1, and Ahnak*. The CD4^+^ *Id2*^+^ cells correspondingly upregulated *S100a6, S100a10, Lgals1, Cxcr3, Cxcr6, Itgb1, and Ahnak*, and in addition upregulated *Lgals3, Ly6a, Ccr2, Capg, Crip1, Ifitm2, Ikzf2* and *Rora*. Transcripts of *Dapl1, Lef1, Lyz2, Igfbp4* and *Plac8* were consistently downregulated in both *CD8*^+^ and *CD4*^+^ *Id2*^+^ T cells **(Figure 1H)**. In both the CD8^+^ and CD4^+^ T cell lineage the *Klrk1*^*+*^ and *Klrc1*^+^ cells showed a highly similar expression, and many upregulated transcripts were also shared with the *Id2*^+^ cells including *Lgals1, Gzma, Gzmb, Zeb2, S100a6, S100a10, Cxcr6, Ahnak, Ccr2*, and *Nkg7*, which underscores the strong relationship between the killer cell lectin receptor (KLR) family, granzymes, and chemokines **(Figure S1C)**. Moreover, downregulation of *Dapl1* and *Igfbp4* was also apparent. All together these data indicate that combination therapy targeting OX40 and PD-L1 promotes expression of molecules associated with cytotoxicity and migration in responding CD8^+^ and CD4^+^ T cell subsets residing in the blood circulation

### Circulating therapy-responsive T cell subsets display effector cell properties with increased cytotoxic and migratory capacity

To validate the association of the *Id2* transcripts with the transcripts of the *Klr* genes in subsets of the *Id2*^*+*^ cells at the protein level, the expression of the KLR family and other effector T cell markers were co-stained with ID2 in circulating T cells obtained from tumor challenged PDOX-treated mice. Substantial subsets within the ID2^+^CD8^+^ T cells expressed NKG2A, NKG2D and KLRG1, whereas ID2^-^ CD8^+^ T cells lacked expression of these KLR family members. ID2^+^CD4^+^ T cells also expressed NKG2A and KLRG1, albeit to a much lesser extent as ID2^+^CD8^+^ T cells. ID2^-^CD4^+^ T cells were devoid of NKG2A, NKG2D or KLRG1 (**Figure 2A**).

**Figure 2.**
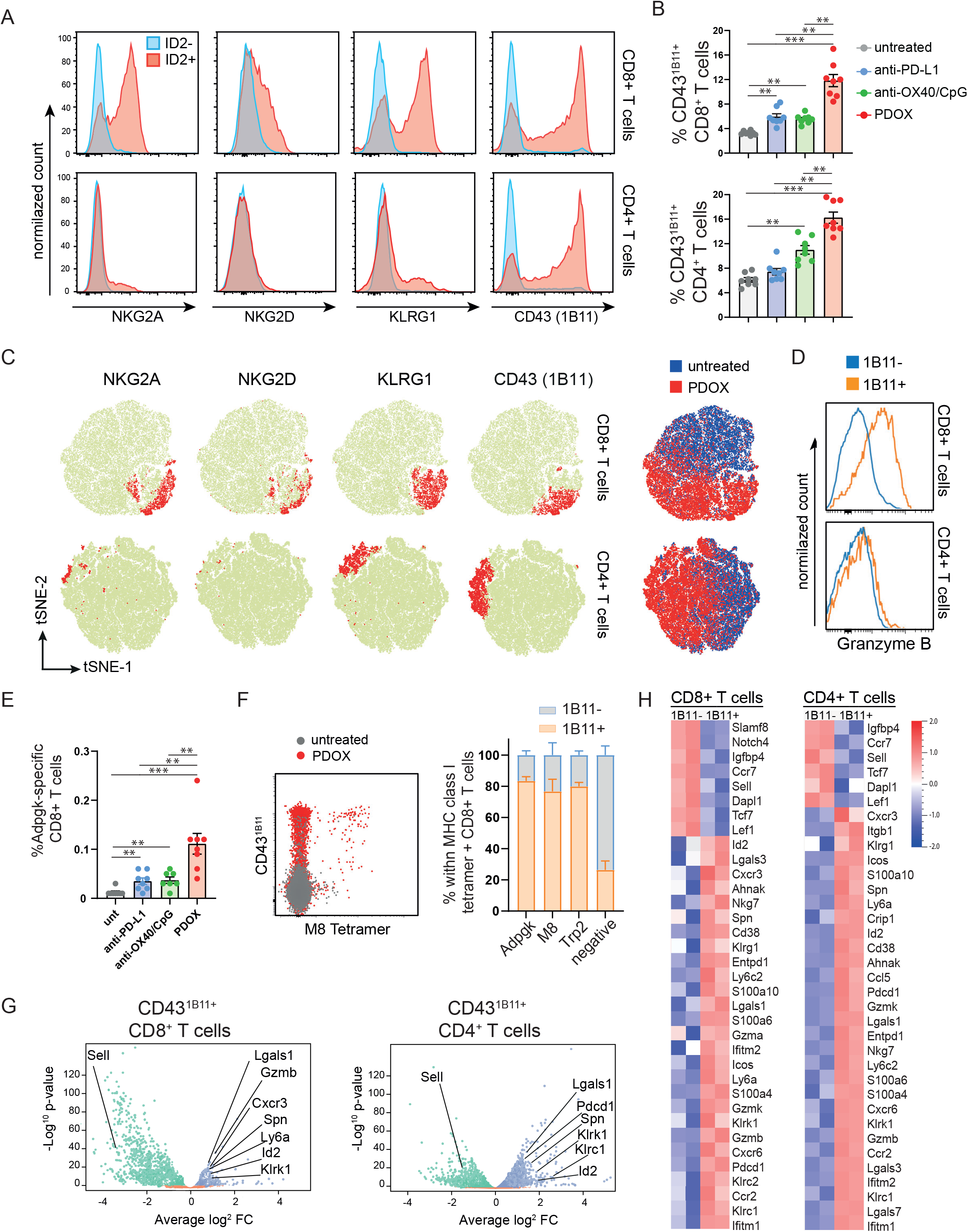
Circulating therapy-responsive T cell subsets display effector cell properties with increased cytotoxic and migratory capacity. (A) Representative histogram plots of NKG2A, NKG2D, KLRG1 and CD43^1B11^ expression on gated ID2^-^ or ID2^+^ CD8^+^ and on gated ID2^-^ or ID2^+^ CD4^+^ T cell populations residing in the blood circulation of MC-38 challenged PDOX treated mice. (B) Percentage of CD43^1B11+^ cells within the total CD8^+^ and CD4^+^ T cell population in blood of untreated and ICT treated groups. Data are represented as mean ± SEM. *p<0.05, **p<0.01, ***p<0.001, by unpaired student’s t test. Each dot represents an individual mouse. (C) tSNE plots of flow cytometric data visualizing NKG2A, NKG2D, KLRG1 and CD43^1B11^ expression (red colored) on gated CD8^+^ and CD4^+^ T cells in the blood from untreated and PDOX treated groups. The blue/red tSNE plot indicates the cell origin to CD8^+^ and CD4^+^ T cells of the untreated and PDOX treated group, respectively. (D) Representative histograms of Granzyme B expression in blood circulating CD43^1B11+^ and CD43^1B11-^ CD8^+^ and CD43^1B11+^ and CD43^1B11-^ CD4^+^ T cell populations of MC-38 challenged PDOX treated mice. (E) Percentage of Adpgk-specific CD8^+^ T cells in blood of untreated and ICT treated groups. Data are represented as mean ± SEM. *p<0.05, **p<0.01, ***p<0.001, by unpaired student’s t test. Each dot represents an individual mouse. Data shown in A-E are representative of two independent experiments. (F) Representative flow cytometry plot of CD43^1B11^ expression on blood circulating M8-specific CD8^+^ T cells. Bar graphs indicate the proportion of CD43^1B11-^ or CD43^1B11+^ cells within the Adpgk, M8, Trp2-specific CD8^+^ T cells and within the M8/Trp2-negative CD8^+^ T cells in blood after MC-38 or HCmel12 tumor challenged mice treated with PDOX. (G) Volcano plots of RNA-seq data of sorted CD43^1B11+^ and CD43^1B11-^ CD8^+^ and CD43^1B11+^ and CD43^1B11-^ CD4^+^ T cells from spleens isolated from wild-type mice challenged with MC-38 and treated with PDOX (day 13 post challenge). (H) Heatmaps (of the data shown in 2G) displaying scaled expression values of discriminating genes.

In order to further characterize the total ID2^+^ subset in both CD4^+^ and CD8^+^ T cells, we examined other surface markers for effector T cells including the O-glycan form of CD43 (sialoforine), which is known as an inducible CD43 isoform expressed transiently by activated T cells (44). This activation-associated isoform is visualized by the 1B11 antibody clone while the antibody clone S1, recognizes CD43 regardless of glycosylation, and reacts with virtually all T cells being activated or not **(Figure S2A)**. Strikingly, the vast majority of the ID2^+^CD8^+^ T cells expressed the hyperglycosylated CD43^1B11^ isoform in contrast to the ID2^-^CD8^+^ T cells. Notably similar results were found for CD4^+^ T cells (**Figure 2A**).

As expected from the connection with ID2 expression, CD43^1B11^-expressing CD8^+^ and CD4^+^ T cells were significantly increased upon PDOX treatment in blood of MC-38 and HCmel12 challenged mice (**Figure 2B, Figure S2B**), and the expression of NKG2A, NKG2D, and KLRG1 showed substantial overlap with CD43^1B11^ expression (**Figure 2C**). Moreover, blood-circulating CD43^1B11^-expressing CD8^+^ but not CD4^+^ T cells from PDOX treated mice expressed high levels of granzyme B (**Figure 2D**). Strikingly, tumor-specific CD8^+^ T cells recognizing the neoantigen Adpgk expressed by MC-38 tumors were increased in the blood circulation **Figure 2E)**, and the vast majority of these cells were positive for CD43^1B11^ **(Figure 2F)**. Likewise, the percentage of circulating CD8^+^ T cells recognizing the tumor antigens Trp2 and M8 in HCmel12 tumor-bearing mice was enlarged after PDOX treatment, and these tumor-specific T cells highly expressed CD43^1B11^ **(Figure 2F)**.

To define and cross-validate the transcriptional signature of the circulating CD43^1B11+^ T cell subsets we performed bulk mRNA sequencing on fluorescence cell sorted CD43^1B11-^ and CD43^1B11+^ CD8^+^ and CD43^1B11-^ and CD43^1B11+^ CD4^+^ T cells from MC-38 tumor-challenged mice that were treated with PDOX. Definitely, as also detected by scRNA-seq both the circulating CD43^1B11+^CD8^+^ and CD43^1B11+^CD4^+^ T cells had analogous upregulated and down-regulated genes as the genes identified in the CD4-T3 and/or CD8-T3 clusters (e.g. *Id2, Lgals1, Klrc1, Klrg1, Klrk1, Ly6a, Cxcr3, Cxcr6, Ccr2, Gzma, Gzmb, Gzmk*) **(Figure 2G, 2H)**. Thus, the circulating therapy-responsive T cell subsets display the induction of a transcriptional program that regulates the cytotoxic and migratory capacity.

### Dynamic induction of therapy-responsive T cell subsets

To gain insight into the dynamics of the therapy-responsive T cell subsets, we longitudinally followed the CD43^1B11+^, the NKG2A^+^ and KLRG1^+^ T cell subsets in the blood circulation of ICT treated MC-38 tumor-bearing mice. Anti-OX40/CpG treatment but not anti-PD-L1 treatment increased the CD43^1B11+^CD8^+^ T cell subset at day 13 post tumor challenge (6 days after the start of the anti-OX40/CpG treatment, and 3 days after the anti-PD-L1 treatment), while at later time-points these treatments resulted in a comparable increase compared to untreated mice. PDOX treatment resulted in a much stronger increase of the CD43^1B11+^CD8^+^ T cells at day 13 compared to anti-OX40/CpG and anti-PD-L1 treatment, and this increase was even more pronounced at day 18 post-treatment **(Figure 3A)**. At day 25 post-tumor challenge, the percentage of CD43^1B11+^CD8^+^ T cells was decreased, yet still higher than anti-OX40/CpG and anti-PD-L1 treated mice. As projected based on the expression profile, the NKG2A^+^ and KLRG1^+^ CD8^+^ T cell subsets showed similar kinetics **(Figure 3A)**. PDOX treatment induced also the highest level of circulating CD43^1B11+^ CD4^+^ T cells at day 13 and day 18 post tumor challenge compared to other treatment groups (anti-OX40/CpG, anti-PD-L1) and untreated mice **(Figure 3B)**. The PDOX-induced CD43^1B11+^ CD4^+^ T cells reached its peak already 13 days after tumor challenge **(Figure 3B)**. Remarkably, the kinetics of CD43^1B11+^ T cells in non-tumor bearing mice was comparable to those in tumor-bearing mice except that PDOX treatment amplified the induction of the CD43^1B11+^ T cells (**Figure 3C**), indicating that the synergy between anti-OX40/CpG and anti-PD-L1 blockade to elicit the highest percentage of CD43^1B11+^ T cells in the blood circulation is also driven by tumor antigens and/or tumor-associated inflammation. A correlation between the level of CD43^1B11+^ CD8^+^ T cells circulating in the blood and MC-38 and HCmel12 tumor survival could be established (**Figure 3D**). Together, these data indicate that the therapy elicited effector T cells, identified by the activation markers CD43^1B11^, NKG2A and KLRG1, are characterized by a dynamic expansion/contraction kinetics resembling that of vaccine or infection provoked T cell responses.

**Figure 3.**
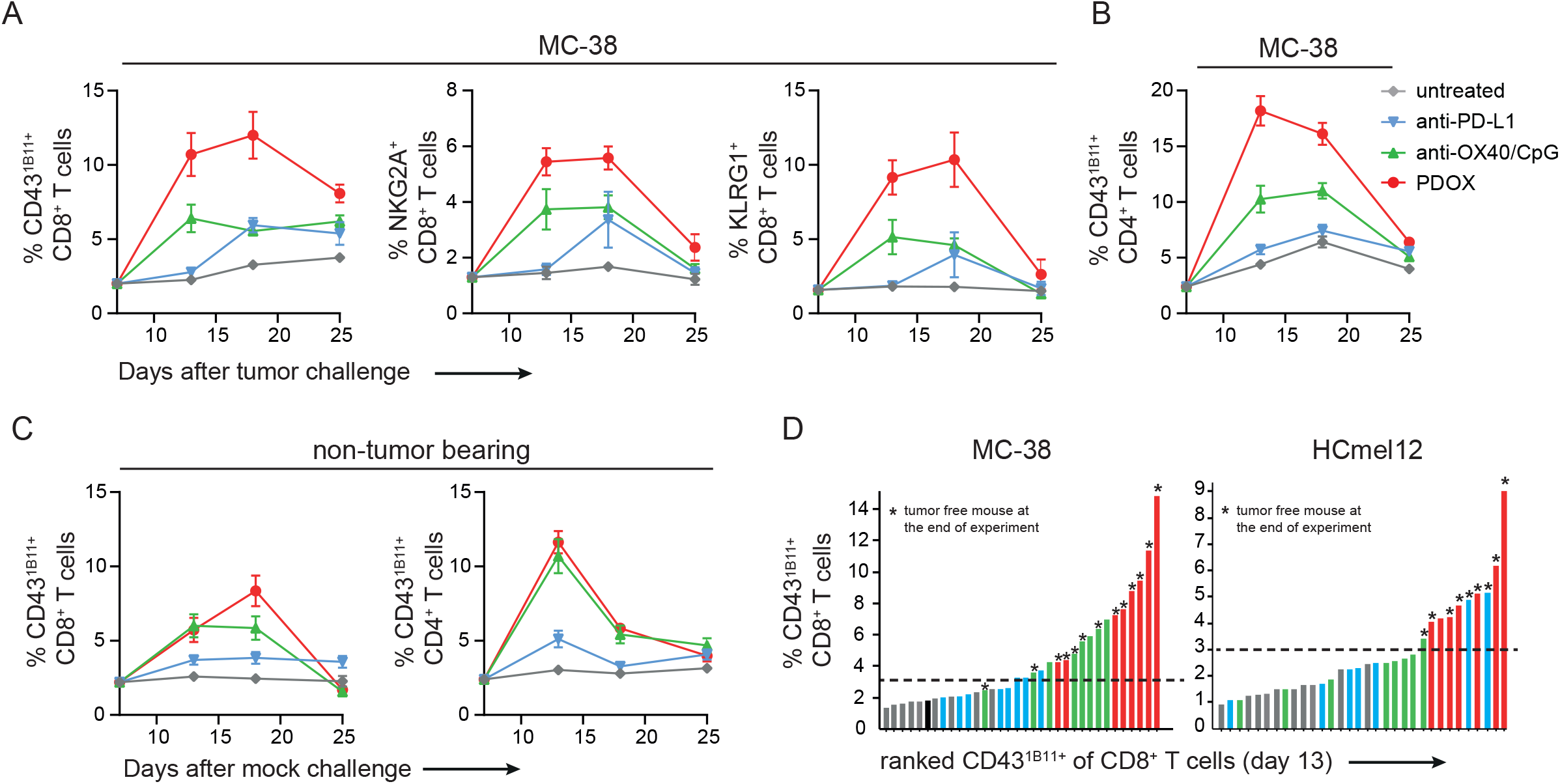
Dynamic induction of therapy-responsive T cell subsets in the blood circulation. (A, B) Kinetics of the CD43^1B11+^, NKG2A^+^, and KLRG1^+^ cells of CD8^+^ T cells (A) and CD43^1B11+^ cells of CD4^+^ T cells (B), in the blood circulation after challenge with MC-38 tumor cells and treated or not with different ICTs (anti-PD-L1, anti-OX40, PDOX). (C) Kinetics of the CD43^1B11+^ cells of CD8^+^ and CD4^+^ T cells after mock challenge and treated similarly as in (A, B). Average frequencies within the total CD8^+^ or CD4^+^ T cell population ± SEM are shown. (D) Ranking of the percentage CD43^1B11+^ cells of CD8^+^ T cells in blood (at day 13) for each individual MC38-bearing-mice (left panel) or HCmel12-bearing-mice (right panel). An asterisk (*) indicates the correlation with tumor-free mice.

### System-wide induction of immunotherapy-responsive T cell subsets

To interrogate whether the PDOX therapy-responsive effector T cell states were elicited system-wide and whether these were interconnected, we assessed the phenotype of the ICT-induced T cell states in several lymphoid tissues at day 18 post MC-38 tumor challenge by CyTOF mass cytometry with 34 cell surface markers, which allowed the identification of T cell signatures in-depth. The marker panel consisted of lineage markers and markers specific for effector T cell activation, differentiation, trafficking and function, and included CD43^1B11^, NKG2A, KLRG1, CXCR3, CD62L, and the ectoenzymes CD38 and CD39 (**Figure 4A, Figure S3**). Viable CD4^+^ and CD8^+^ T cells were manually gated from all acquired events and subsequently analyzed by HSNE using Cytosplore (45) and Cytofast (46, 47). We selected clusters based on the significant difference (p<0.05) and average abundance (>5%). Remarkably, in the blood compartment, all CD8^+^ and CD4^+^ T cell clusters that were significantly higher in PDOX treated mice expressed CD43^1B11^ (**Figure 4B, Figure S4**). Two of the four CD43^1B11+^CD8^+^ T cell clusters were most abundant in the PDOX treated group compared to all other groups and co-expressed either KLRG1, CD38, CD39, PD-1, LAG-3 (Blood cluster CD8-5) or co-expressed the same markers and in addition NKG2A and ICOS (Blood cluster CD8-11). Three CD43^1B11+^CD4 T cell clusters, which were more abundant in the PDOX treated group compared to all other groups, all co-expressed CXCR3 and ICOS (CD278), and differentially expressed PD-1, CD38, CD39. In the spleen, one CD8^+^ T cell cluster expressing CD43^1B11^ (Spleen CD8-13) was most abundant in PDOX treated mice, and similar to blood cluster CD8-5 these cells co-expressed KLRG1, CD38, CD39, PD-1, and LAG-3. Four splenic CD4^+^ T cell clusters expressing CD43^1B11^ were higher in PDOX treated mice and highly similar to the CD43^1B11^ expressing CD4^+^ T cells in blood. In the lymph node and bone marrow significantly higher frequencies of CD43^1B11+^CD4^+^ T cells but not CD43^1B11+^CD8^+^ T cells in the PDOX treated mice were detected, and these cells differentially expressed ICOS, CD38, CD39, and KLRG1. Thus, besides residing in the blood circulation also in other lymphoid compartments CD8^+^ and CD4^+^ T cell clusters expressing CD43^1B11^ and co-expressing NK cell receptors, chemokine receptors and ectoenzymes are elicited that connect to effective immunotherapy.

**Figure 4.**
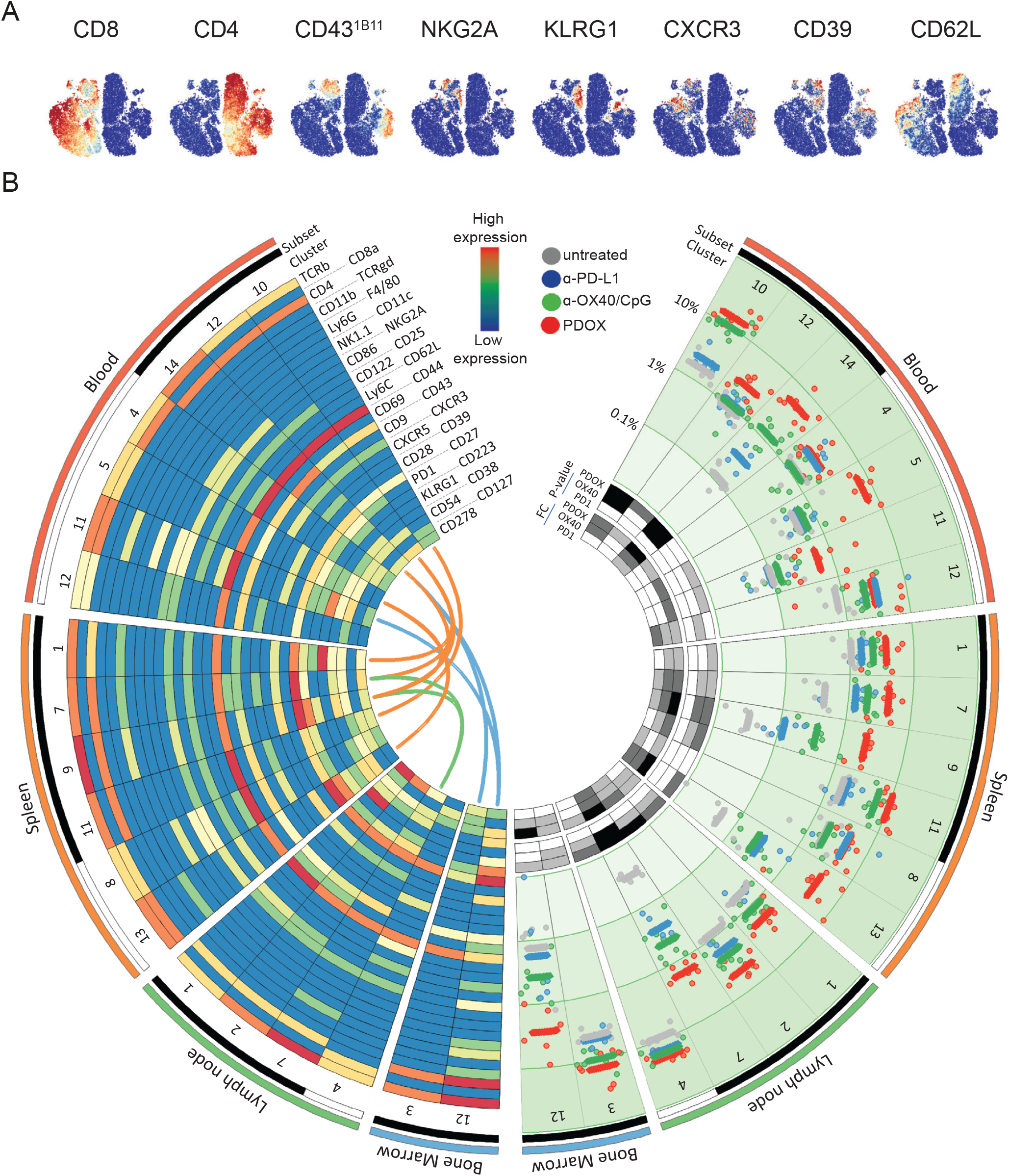
Systemic induction of therapy-responsive T cell subsets upon effective ICT. (A) tSNE plots of blood T cells isolated from tumor-bearing untreated and ICT treated (anti-PD-L1, anti-OX40/CpG, PDOX) mice visualizing the expression intensity of cell-surface markers measured by CyTOF mass cytometry. (B) Circle diagram showing significantly different T cell clusters in blood, spleen, lymph nodes and bone marrow measured by CyTOF. Clusters are mirrored with phenotypic information on the left and abundance on the right. From outside in: tissues are depicted in coloured bars, CD8^+^ and CD4^+^ T cell subsets as white and black bars, respectively. Cell clusters are indicated with numbers. On the left, a heatmap for 31 relevant markers depicts normalized expression for each marker per cluster. Clusters that are highly correlated (r>0.7) between compartments are connected by coloured lines. On the right, frequency of subsets within its parent is shown per cluster on a log-scale. Each dot represents an individual mouse. Mean values per group are indicated by lines. Multiple-testing corrected p-values for each of the ICT treated groups compared to the untreated group are shown, with p>0.01 (white), p<0.01 (light grey), p<0.001 (dark grey), p<0.0001 (black). P-values were based on t-test on log-transformed frequencies and corrected by Benjamini-Hochberg correction. Means of fold change (FC) of log-transformed values of ICT groups over the untreated group are shown with FC<2 (white), FC>2 (light grey), FC>4 (dark grey), FC>16 (black).

To determine the correlation between the identified therapy-responsive T cell clusters across all lymphoid tissues via an unbiased approached, Spearman’s correlation analyses were performed system-wide **(Figure 4B)**. The therapy-responsive CD4^+^ and CD8^+^ T cell clusters in the blood are closely related to those in the spleen (*r* > 0.70), and blood CD4^+^ T cell clusters are also connected to the bone marrow CD4^+^ T cell clusters. Moreover, lymph node CD4^+^ T cell cluster 7 relates to splenic CD4^+^ T cell clusters. Together, these data show an inter-connectivity between therapy-responsive T cell clusters residing in different lymphoid tissues, indicating the induction of system-wide effects of ICT enabling efficient tumor immunity.

### Expansion of the therapy-responsive CD8 T cell subset in the blood circulation and tumor-microenvironment is modulated by NK cell receptors

We next analyzed the therapy-responsive T cells in the TME of MC-38 challenged mice. Because PDOX treatment is very effective, the treatment started at later time-points (around day 10) to obtain sufficient tumor material for analysis. Compared to untreated animals, PDOX treatment increased the percentage of leukocytes in the TME, which was mainly caused by an increase in CD8^+^ T cells, and this correlated with an increase in tumor-specific CD8^+^ T cells (**Figure 5A**). In the TME of HCmel12-challenged mice similar data was obtained (**Figure 5B**). Detection of CD8^+^ T cells by immunofluorescent staining showed that PDOX treatment promoted higher numbers and deep infiltration of CD8^+^ T cells into the TME (**Figure 5C**).

**Figure 5.**
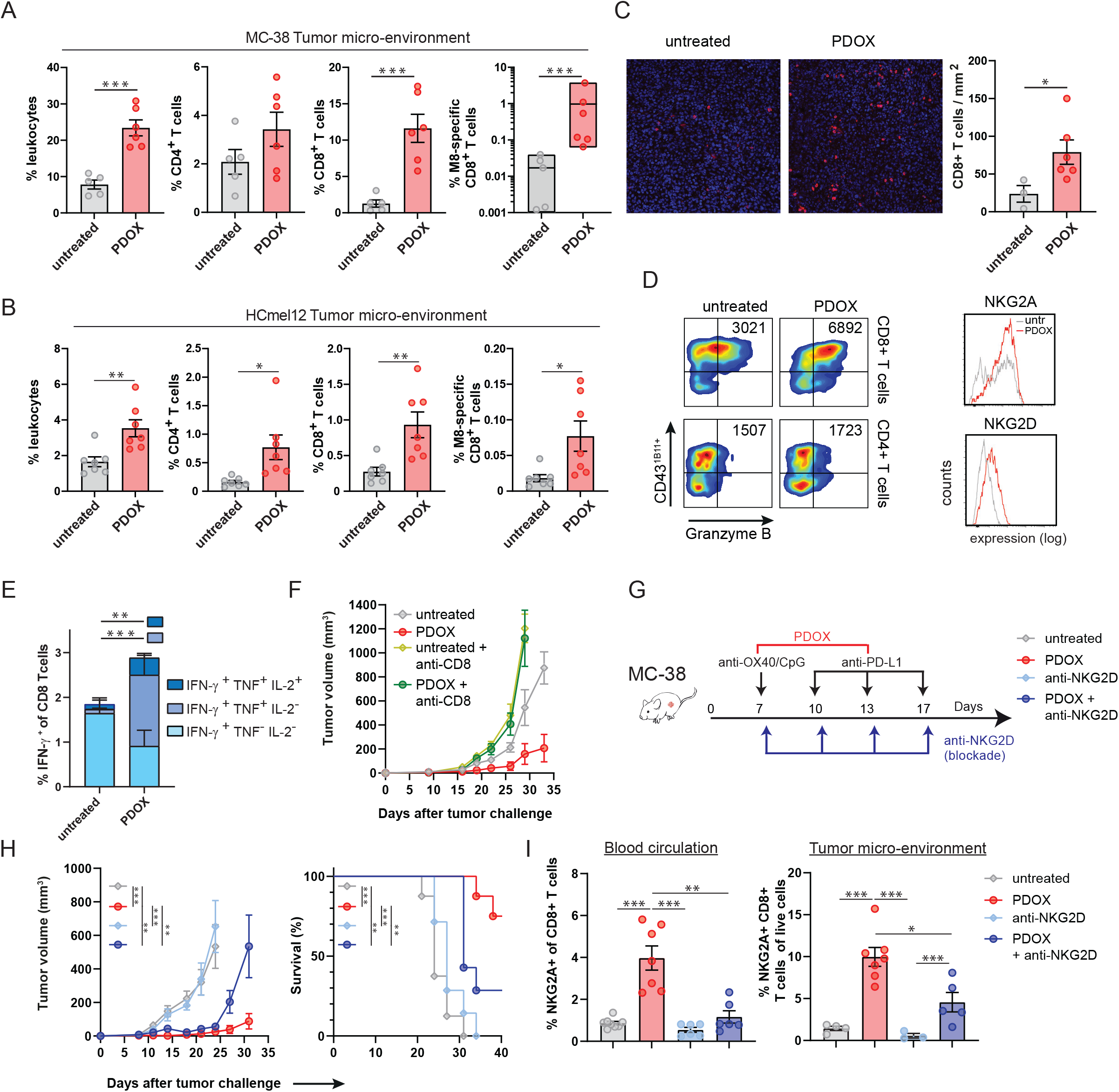
Expansion of the therapy-responsive CD8 T cell subset in the blood circulation and tumor-microenvironment is modulated by NK cell receptors. (A-B) Percentage leukocytes, total CD8^+^ and CD4^+^ T cells of live cells and percentage M8-specific CD8^+^ T cells in the MC-38 (A) and HCmel12 tumor micro-environment of untreated and PDOX treated mice. Data are represented as mean ± SEM. Each dot represents an individual mouse. (C) Representative immunofluorescence images of MC-38 tumor-infiltrated CD8^+^ T cells (red colored) of untreated and PDOX-treated mice. Bar graph indicates the absolute CD8^+^ T cell count per mm^2^. (D) Representative flow cytometry plots indicate CD43^1B11^ versus granzyme B expression of tumor-infiltrated CD8^+^ and CD4^+^ T cells. Granzyme B expression numbers indicate the average intensity of Granzyme of CD43^1B11+^ T cells. Representative histograms show NKG2A and NKG2D expression on MC-38 tumor-infiltrated CD8^+^ T cells of PDOX treated and untreated mice. (E) The proportion of single, double and triple cytokine producing cells within the tumor-infiltrating CD8^+^ T cells of MC-38 tumor challenged mice treated or not with PDOX. (F) Average MC-38 tumor growth of untreated and PDOX treated wild-type mice receiving CD8 depleting antibodies or mock. (G) Schematic of the strategy. C57BL/6 mice were subcutaneously challenged with MC-38 syngenic tumors and were left untreated or treated with PDOX in combination with blocking NKG2D antibodies. (H) MC-38 tumor growth (mean ± SEM) and survival curves of untreated and PDOX treated mice in combination with blocking anti-NKG2D antibodies. One of 2 independent experiments is shown. (I) Percentage NKG2A^+^ cells of blood CD8^+^ T cells and percentage NKG2A^+^CD8^+^ T cells of live cells in the tumor-micro-environment of untreated and PDOX treated mice in combination with blocking NKG2D antibodies. *p<0.05, **p<0.01, ***p<0.001

To assess the effector potential of the tumor-infiltrated T cells, granzyme and cytokine production was evaluated. In both untreated and PDOX treated mice the vast majority of tumor-infiltrated CD8^+^ and CD4^+^ T cells expressed CD43^1B11^ (**Figure 5D**). However, upon PDOX treatment the granzyme B expression was increased in the tumor-infiltrated CD8^+^ T cells, and this correlated with a higher NKG2A and NKG2D expression (**Figure 5D**). In addition, PDOX treatment elicited high percentages of polyfunctional cytokine producing CD8^+^ T cells, i.e. double producing IFN-γ + TNF and triple producing IFN-γ + TNF + IL-2, whereas the majority of tumor-infiltrated CD8^+^ T cells in the untreated group were single IFN-γ producers (**Figure 5E**). In line with the increment of cytotoxic and cytokine polyfunctional tumor-infiltrated CD8^+^ T cells, the depletion of CD8^+^ T cells completely dismantled the efficacy of PDOX treatment (**Figure 5F**).

To functionally assess the NK cell receptor expression on CD8^+^ T cells, we used antibodies that blocked NKG2D (**Figure 5G**), which is known as a molecule with the capacity to provide costimulatory signals to T cells (48). Blockade of NKG2D reduced the effectivity of the PDOX treatment on controlling tumor outgrowth, whereas NKG2D blockade in untreated mice had no significance (**Figure 5H**). This effect of NKG2D blockade on PDOX treatment related to a diminution of NKG2A^+^ CD8^+^ T cells in the blood circulation and TME **(Figure 5H)**. We conclude that NK cell receptor expressing subsets are instrumental for the PDOX therapeutic efficacy with NKG2D supporting the systemic stimulation of the therapy-responsive effector CD8^+^ T cells.

### CD43 and CXCR3 expression supports the tumor migration of therapy-responsive T cell subsets

To functionally assess whether the CD43 expression is critical for the efficacy of ICT we examined the PDOX-responsiveness in settings of CD43 availability and absence. For this, wild-type mice and mice deficient in the *Spn* gene (coding for CD43) were challenged with MC-38 tumor cells, and left untreated or treated with PDOX (**Figure 6A, 6B**). Whereas CD43 proficient mice showed the anticipated therapeutic efficacy of PDOX treatment upon tumor challenge, mice deficient in CD43 could not control MC-38 tumor outgrowth despite PDOX treatment, (**Figure 6C**). Remarkably, CD43 deficiency did not affect the number of NKG2A^+^ CD8^+^ T cells in the blood but rather resulted in a diminution of NKG2A^+^ CD8^+^ T cells in the TME (**Figure 6D**), indicating that the tumor migration of therapy-responsive T cells is regulated by CD43, which is in line with the ability of the CD43^1B11^ isoform to function as a ligand for the cell adhesion molecule E-selectin (49).

**Figure 6.**
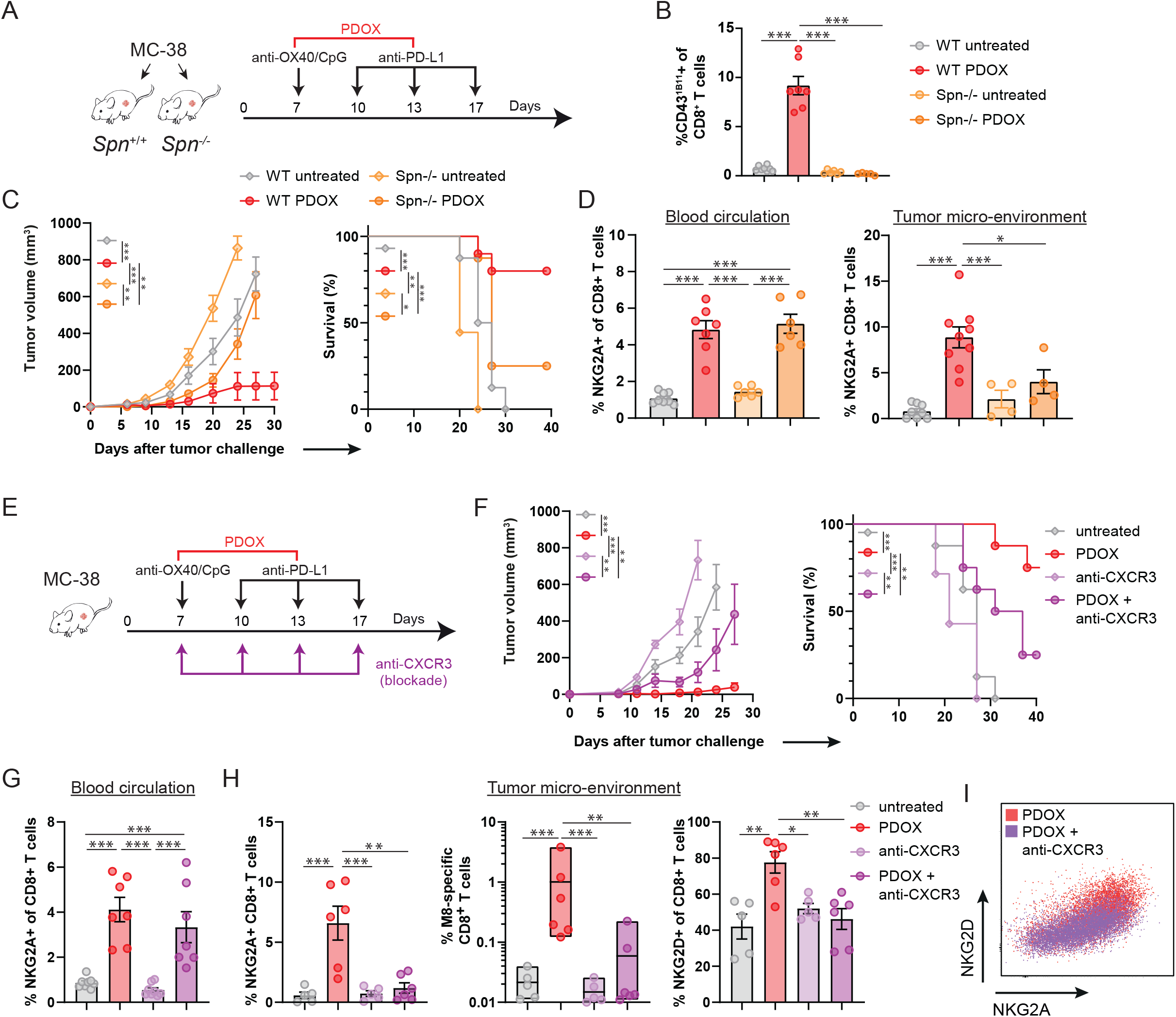
CD43 and CXCR3 expression supports the tumor migration of therapy-responsive T cell subsets. (A) Schematic of the strategy. C57BL/6 (WT) and Spn^-/-^ mice were subcutaneously challenged with MC-38 tumors and were left untreated or treated with PDOX. (B) Percentage CD43^1B11+^ cells of blood CD8^+^ T cells untreated and PDOX treated WT and Spn^-/-^ mice mice. (C) MC-38 tumor growth (mean ± SEM) and survival curves of untreated and PDOX treated WT and Spn^-/-^ mice. One of 2 independent experiments is shown. (D) Percentage NKG2A^+^ cells of blood CD8^+^ T cells and percentage NKG2A^+^CD8^+^ T cells of live cells in the TME of untreated and PDOX treated WT and Spn^-/-^ mice. (E) Schematic of the strategy. WT mice were subcutaneously challenged with MC-38 syngenic tumors and were left untreated or treated with PDOX in combination with blocking CXCR3 antibodies. (F) MC-38 tumor growth (mean ± SEM) and survival curves of untreated and PDOX treated mice in combination with CXCR3 blocking antibodies. One of 2 independent experiments is shown. (G) Percentage NKG2A^+^ cells of CD8^+^ T cells in blood of untreated and PDOX treated mice in combination with blocking CXCR3 antibodies. (H) Percentage NKG2A^+^ CD8^+^ T cells of live cells, percentage M8-specific CD8^+^ T cells and percentage NKG2D^+^ cells of CD8^+^ T cells in the tumor-micro-environment of untreated and PDOX treated mice in combination with blocking CXCR3 antibodies. (I) Representative flow cytometry plot of NKG2A *versus* NKG2D expression of tumor-infiltrated CD8^+^ T cells in PDOX treated mice in combination with blocking CXCR3 antibodies. *p<0.05, **p<0.01, ***p<0.001

To test whether the chemokine receptor CXCR3, expressed on the CD43^1B11+^ T cell subset and known to mediate adhesion-induction (50) was also implicated in tumor migration we blocked this chemokine receptor by antibodies provided during PDOX treatment of tumor challenged mice (**Figure 6E**). Obstruction of CXCR3 resulted in reduced effectivity of PDOX to control tumor outgrowth (**Figure 6F**). This effect of CXCR3 blockade related to decreased infiltration of CD43^1B11+^ CD8^+^ T cells in the TME as well as reduced tumor-infiltrated NKG2A^+^ CD8^+^ T cells and tumor-specific CD8^+^ T cells, while the CD43^1B11+^ and NKG2A^+^ CD8^+^ T cells were unaffected in the blood circulation **(Figure 6G and 6H, Figure S6)**. Moreover, CXCR3 blockade specifically prevented NKG2D^+^ CD8^+^ T cells into to the TME **(Figure 6H, 6I)**. Altogether, we conclude that CD43 and CXCR3 mediate the tumor migration of the therapy-responsive CD8^+^ T cells, which is critical for these cells to achieve tumor control.

### Identification of NK cell receptor expressing CD8^+^ T cell subsets in the blood circulation of PD-1 therapy-responsive patients

To determine whether corresponding therapy-responsive T cell subsets could be identified in the blood circulation of patients receiving ICT, we evaluated anti-PD-1 responding and non-responding patients with melanoma or non-small cell lung cancer (NSCLC) (i.e. 9 melanoma (5 responders and 4 non-responders), 5 NSCLC (3 responders and 2 non-responders). Peripheral blood was collected 2 weeks after treatment and stained with a panel of antibodies that incorporated the detection of NK cell receptors expressed by activated human CD8^+^ T cells, i.e., KLRG1, CD56 and KLRB1. (**Figure S4**). Samples were subjected to CyTOF mass cytometry and data analysis by HSNE and Cytofast revealed two distinct CD8^+^ T cell subsets that were significantly increased in the anti-PD-1 responding group (CD8-24 and CD8-17) as compared to the anti-PD-1 non-responders **(Figure 7A, 7B, 7C)**. Both CD8-24 and CD8-17 subsets expressed KLRG1, CD38 and CD29, and exclusively expressed CD56 and KLRB1.

**Figure 7.**
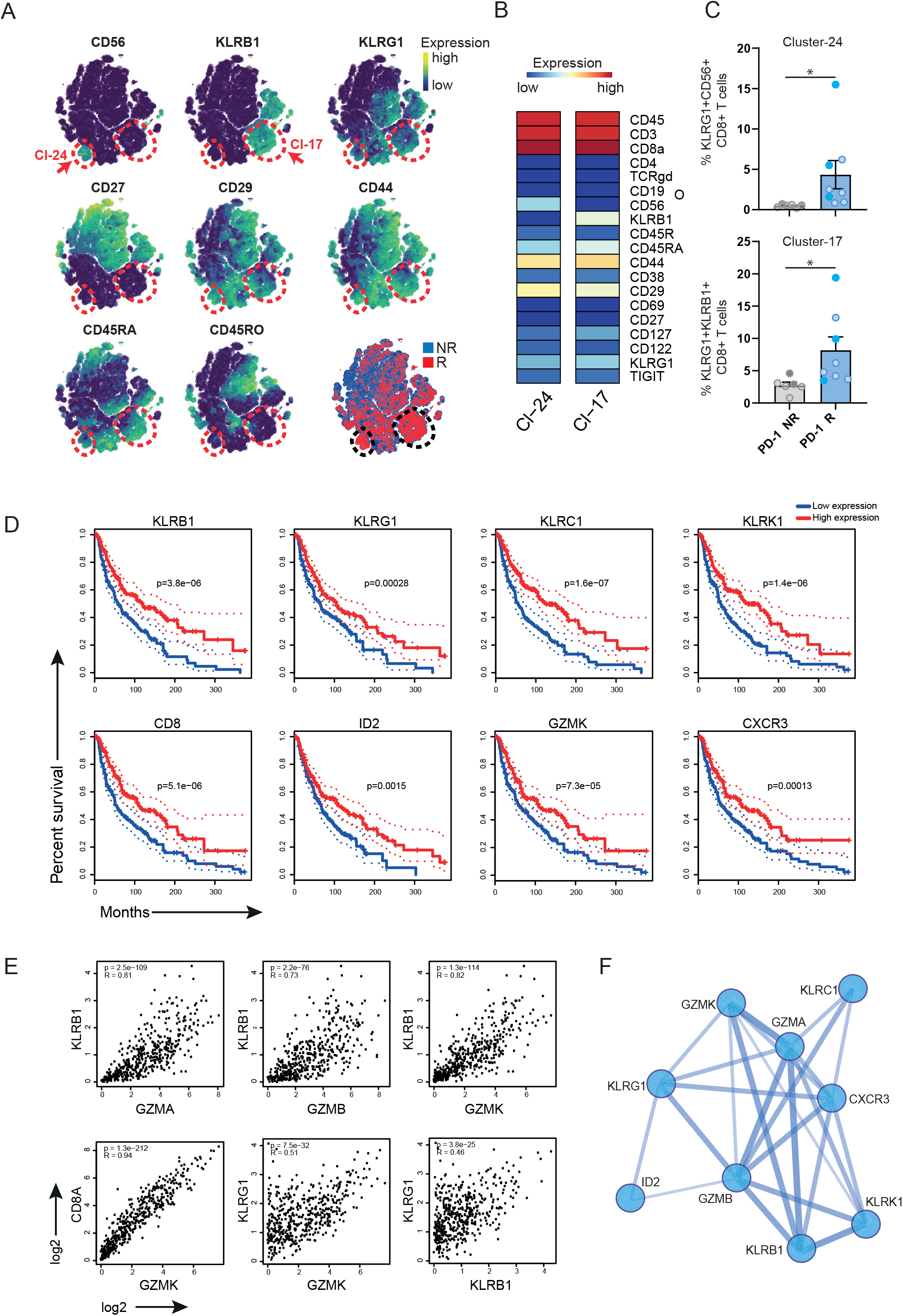
Identification of NK cell receptor expressing CD8^+^ T cell subsets in the blood circulation of PD-1 therapy-responsive patients. (A) tSNE embedding showing the level of expression marker on the blood circulating CD8^+^ T cell population of PD-1 (non-)responding patients after 2 weeks therapy. The arrows indicate cluster 24 and 17. Data shown is pooled from the responder and non-responder groups of melanoma and lung cancer patients. Level of marker expression is displayed by a scale from blue to yellow. Lower right tSNE plot indicates cells color coded per responder (red) and non-responder (blue). (B) Heatmaps of CD8^+^ T cell clusters 24 and 17. Level of ArcSinh5-transformed expression marker is displayed by a rainbow scale. (C) Percentage of KLRG1^+^CD56^+^ (cluster 24) and KLRG1^+^KLRB1^+^ (cluster 17) of the blood circulating CD8^+^ T cell population of PD-1 (non-)responding patients. Bar graphs represent average frequency (+ SEM). Circles represent samples of non-responder melanoma (light grey), responder melanoma (light blue), non-responder lung cancer (grey) and responder (blue) patients. *p<0.05 (D) The Cancer Genome Atlas (TCGA) survival plots for high *versus* low marker gene expression for skin cutaneous melanoma (SKCM). (E) TCGA correlation analysis of therapy-responsive marker genes for SKCM. Spearman correlation coefficient is indicated. (F) Protein network analysis of the markers expressed by therapy-responsive T cells. Line thickness indicates the strength of support for interaction.

Next, we evaluated the survival and correlation of the therapy-responsive marker genes discovered in the mouse and human studies in large cohorts of patients with skin cutaneous melanoma. Higher expression of the KLR family members (KLRB1, KLRG1, KLRC1 and KLRK1), granzymes as well as CXCR3 and ID2 associated with better prognosis (**Figure 7D**). Correlation analysis of these genes indicated a strong association of the granzyme family with the KLR family members and with CD8 (**Figure 7E**). Moreover, non-biased protein interaction analysis confirmed the connection between the therapy-responsive markers indicating an underlying transcriptional program (**Figure 7F**). Thus, comparable to the findings in experimental settings, effective ICT in patients correlates to increment of CD8^+^ T cell subsets characterized by programming for cytotoxic effector function.

## Discussion

A major challenge for ICT is to overcome the substantial variability of this therapy through the identification of predictive biomarkers. Ideally, such biomarkers exist in easily accessible compartments, such as the blood circulation, and also reflect the therapy response in the TME. Here, we demonstrated in two different murine tumor models, MC-38 (colorectal carcinoma) and HCmel12 (melanoma), that a dynamic and systemic T cell response develops upon efficient immunotherapy, which is characterized by an interconnected gene signature related to cytotoxicity and migration. Furthermore, analysis of peripheral blood samples of PD-1 blockade therapy-responsive and unresponsive patients, and of gene expression profiles in melanoma highlight the potential utility of the cytotoxic gene signature consisting of several NK cell markers expressed by CD8^+^ T cells.

The combination of an PD-1/PD-L1 pathway antagonist and an OX40 agonist combined with CpG was remarkably efficient in eradicating developing tumors. These data show the possible benefit of using combinatorial treatment of already used therapeutics in patients, e.g. anti-OX40 (51) and anti-PD-L1 (52), and emphasized the potential of this combination as has also been observed in other mouse tumor models (53, 54). This harmonizing effect of the combinatorial treatment was deciphered by complementary high-dimensional single-cell technology platforms. Both scRNA-seq and mass cytometry highlighted functionally active CD4^+^ and CD8^+^ T cell states that were dynamic in their kinetics and characterized by NK cell receptor expression and by expression of adhesion/migration receptors. The kinetics of the therapy-responsive T cells, i.e. sharp expansion followed by a contraction phase, which is typical for acute infection, may reflect a temporal systemic activation but the T cell activation may be ongoing in the TME. Albeit, less impressive, this expansion was also observed following OX40/CpG treatment while PD-L1 blockade seems to mainly facilitate this T cell expansion when combined. Additional PD-1 upregulation occurring following OX40 triggering may thus be efficiently counter-acted, and allows the T cells to rapidly expand and differentiate to cytotoxic effector T cells. Albeit the expression of OX40 on CD4^+^ T cells is higher as compared to CD8^+^ T cells, this mechanism is likely to occur in both CD4^+^ and CD8^+^ T cells because direct triggering of OX40 on either CD4+ and CD8^+^ T cells results in effector T cell formation (55, 56). Nevertheless, the OX40-activated CD4^+^ T cells may additionally help the CD8^+^ T cells in their expansion and differentiation (57). Whether triggering of related receptors, such as 4-1BB or CD27 belonging also to the TNFR superfamily, has similar effects as described here for OX40 remains to determined.

Our study also indicated that the efficiency of the immunotherapeutic treatment is mirrored by the induction of peripheral T cells that could be identified by cell surface markers. The cell surface expression of the hyperglycosylated form of CD43, and the NK cell receptors NKG2A, NKG2D and KLRG1 on CD8^+^ T cells was a strong signature for these cells as an indicator for cytotoxic effector function based on the co-expression with granzymes. In line with this are findings in the TME of melanoma patients, where KLRG1 was found to be expressed on the cytotoxic T cell compartment (22). Moreover, fate-tracking studies in mice indicated that the KLRG1^+^ CD8^+^ T cells display developmental plasticity and basically these cells can differentiate into all memory T cell lineages, which underscores the value of this marker (58). Indeed, KLRG1^+^ CD8^+^ T cells are excellent predictors of the effectivity of cancer vaccines (59). In the PBMC compartment of patients we also observed KLRB1 (CD161), which was recently discovered as functional marker on human CD8^+^ T cells in the TME of glioma (60). Besides, effects on CD8^+^ T cells we also observed system-wide expansion and contraction of CD4^+^ T cells expressing CD43^1B11^, KLRG1 and CX3CR3, which is in line with studies on human peripheral blood of anti-PD-1-treated patients in which CXCR3^+^ CD4^+^ T cells correlated with positive clinical outcome (61). Remarkably, transcripts of encoding NK cell receptors (*Klrc1, Klrk1, Klrg1*) were observed in CD4^+^ T cells, however at the protein level only KLRG1 was found to be highly expressed suggesting differential posttranscriptional regulation. Whether NKG2A and NKG2D, which likely have opposing functions with respect to T cell activation, are both concurrently functional is complex given that the ligands of these receptors, i.e. HLA-E, and MICA/B, ULBP1-6, respectively, can be inducibly expressed in both healthy and cancer tissue (62, 63).

The blocking studies targeting the NK cell receptor NKG2D and the chemokine receptor CXCR3, together with the CD43 deficient setting, identified the importance of simultaneous induction of both cytotoxic and migratory properties. Although the combination therapy we used was already effective, blockade of NKG2A in more resistant tumors, which is known to synergize with PD-1 blockade (64), may further improve anti-tumoral responses. Another upregulated molecule upon PDOX treatment that could be targeted to improve the outcome is the costimulatory molecule ICOS, which is linked to enhancing PD-1 targeted immunotherapy in mice (65) but also in responsive patients (66).

Additional clinical studies are required to determine the predictive significance of our findings. Studies with agonistic antibodies targeting costimulatory receptors such as OX40 in combination with inhibitory immune checkpoint blockade may be of particular interest. Recent studies already indicated the correlation between effector/effector-memory CD8^+^ T cell responses in the peripheral blood and clinical responses to immune checkpoint blockade (29, 32, 33). Our work highlights these studies and additionally proposes that effective combinatorial therapy is more powerful in the induction of such peripheral T cell responses. Moreover, we show that NK cell receptors and the chemokine receptor CXCR3 are functional markers, and the targeting thereof may further improve the clinical response.

In conclusion, we have provided evidence for an effective-related-treatment immune signature. Future studies entailing a systematic and multicenter cohort of patients with different cancer types for which a combinatorial anti-TNFR family member with anti-PD-1 treatment is approved remains needed. A prediction signature might then be directly used in clinical practice to stratify different levels of effectiveness of treatments.

## Acknowledgements

The authors acknowledge Camilla Labrie and Iris Pardieck for their technical support in performing animal experiments and sample processing, Jannie Borst and Thorbald van Hall for critically reviewing the manuscript, and the Flow cytometry Core Facility of Leiden University Medical Center (LUMC) for their technical support in sample acquisition.

## Author contributions

GB and TvdS designed and performed experiments, and analyzed data. EvdG, BB, TW, SvD, AR, EBN, and MC performed experiments. SJ, TA and FvH, performed computational data analysis. AB, JH and JA provided essential materials. ML and PH generated the CD43 knockout mice. FO provided scientific input and supervision. RA designed the experiments, analyzed data, initiated and supervised the project. GB and RA wrote the manuscript, with all authors contributing to writing and/or providing feedback. All authors read and approved the manuscript.

## Materials and Methods

### Mice

C57BL/6 female mice were obtained from Janvier Laboratories (Le Genest-Saint-Isle, France). At the start of the experiments, mice were 6 to 8 weeks old. Mice were housed in individually ventilated cages (IVC) under specific pathogen-free (SPF) conditions in the animal facility of Leiden University Medical Center (LUMC, Leiden, The Netherlands). All animal experiments were approved by the local and national committees of animal experiments under the permit number AVD116002015271, and performed according to the recommendations and guidelines set by the LUMC and by the Dutch Act on Animal Experimentation and EU Directive 2010/63/EU. Dutch Experiments on Animals Act.

### Generation of *Spn (Cd43)* knockout mice

Spn^-/-^ mice (*Spn*^em1Lumc^; MGI:6360988) were generated using CRISPR-Cas9-mediated targeting of zygotes, which resulted in deletion of the coding sequence of Exon 2 of the *Spn* (*Cd43*) gene. C57BL/6J zygotes were microinjected at embryonic day 2 (E1.5, 2cell stage), with CRISPR/Cas9 RNP complexes with guide sequences CCTCAATCTCTATGAGCAAC and GGTGCAAGGCCATCTCCAGA and transferred to pseudo-pregnant recipients (crRNA, tracrRNA and Cas9 obtained from IDT). Mosaic candidates were selected based on PCR with primers upstream 5’ Crispr and downstream of the 3’ Crispr, followed by Sanger sequencing of the PCR product. One founder animal was selected with a deletion of Chr7: 126,735,256-126,736,568 (GRCm39) encompassing the complete coding sequence of the gene. The line was maintained on a C57BL/J6 background. PCR and sanger sequencing analysis of 5 most likely off-target sites based on CRISPOR.tefor.net scores, showed no Off-targets events (data not shown).

### Cancer Patient Samples

PBMC samples from melanoma and non-small cell lung cancer (NSCLC) patients treated with PD-1 antibodies (pembrolizumab or nivolumab) were collected at the Amsterdam University Medical Center, the Netherlands Cancer Institute and Erasmus MC. PBMCs were isolated and cryopreserved in medium after Ficoll density gradient centrifugation. Ethical approval was provided by the local medical ethical committees, and written informed consent was obtained in accordance with the Declaration of Helsinki.

### Tumor challenge models

Treatment schedule of experiments are indicated in the respective figures and legends. In tumor challenge experiments, mice were inoculated in the flank with 0.3 × 10^6^ MC-38 (subcutaneously, 200 μL volume) or with HCmel12 (intra-dermally, 30 μL volume) tumor cells in PBS containing 0.2% BSA. Tumor outgrowth was monitored by caliper-based measurements in three dimensions. Mice were euthanized when tumor size reached >1000 mm^3^ in volume or when mice lost > 20% of their total body weight (relative to initial body mass).

### Antibody-based interventions in vivo

Antibodies targeting CD8 (clone 2.43), OX40 (clone OX86) and PD-L1 (clone MIH-5) were purified from hybridoma cultures. The agonistic OX40 antibodies were injected subcutaneously along with CpG (ODN1826). The blocking anti-PD-L1 antibodies were administered intraperitoneally. CD8^+^ T cell depleting antibodies were administered intraperitoneally twice weekly (first injection 150 μg/mouse followed by 50 μg/mouse) for 2 weeks starting 4 days before tumor challenge. CXCR3 (clone CXCR3-173), and NKG2D (clone CX5) antibodies were purchased from Bio X Cell (Lebanon, NH, USA) and administered intraperitoneally. Depletion and blockade of the antibodies was verified by flow cytometry.

### Preparation of single-cell suspensions for cytometry

Peripheral blood was collected from the tail vein. Spleen and lymph nodes were minced through 70 µm cell strainers. Bone marrow cells were extracted from the femurs and tibias by flushing with medium containing 8% FBS. Erythrocytes were removed from blood and tissues using a hypotonic ammonium chloride lysis buffer for 2 minutes. Tumors were obtained after transcardial perfusion with 30 mL of PBS/EDTA (2 mM), and after mincing incubated with 2.5 mg/mL Collagenase D and DNAse (Roche) for 20 minutes at 37°C. Single-cell suspensions were obtained by using 70 µm cell strainers (BD Biosciences). All samples were washed with medium containing 8% FBS before further processing.

### Flow cytometry

After the single-cell suspension preparation, samples were washed with staining buffer (PBS with 2% FBS). Mouse Fc-Receptors were blocked with anti-mouse CD16/32 (clone 2.4G2) and 10% naÏve mouse serum for 15 minutes before antibody staining. Following a wash step with PBS, live/dead staining was performed using Zombie NIR (Biolegend) for 10 minutes at room temperature. After washing in staining buffer, cells were stained using combinations of fluorescently labelled antibodies and tetramers for at least 0.5h. MHC class I tetramers (Adpgk – H-2D^b^ tetramer containing the ASMTNMELM peptide, M8 – H-2K^b^ tetramer containing the KSPWFTTL MuLV p15E peptide, Trp2 - H-2K^b^ tetramer containing the SVYDFFVWL TRP2 peptide were all made in house and either labelled with APC or PE. Intracellular Granzyme B and ID2 stainings were performed after fixation in True-Nuclear fixation buffer for 45 min at room temperature and permeabilization in True-Nuclear permeabilization buffer. Intracellular cytokine staining was performed after incubation for 5 hours with Remel™ PHA Purified (ThermoFisher Scientific) in the presence of 2 µg/ml Brefeldin. Samples were acquired on a BD FACS LSR Fortessa X-20 4L (BD Biosciences, San Jose, CA, USA) or a 3L Cytek Aurora flow cytometer at the Flow cytometry Core Facility of Leiden University Medical Center. The data was analyzed using FlowJo v10 (Treestar) software and OMIQ data analysis software.

### CyTOF mass cytometry

After the single-cell suspension preparation, debris and aggregates were removed using a 100/60/40/30-percent gradient of Percoll (GE HealthCare) in RPMI 1640 (Lonza), and pelleted single cells in the 40% fraction were resuspended. Approximately 3 × 10^6^ cells were stained for each sample.

Metal-conjugated antibodies were either purchased from Fluidigm Sciences or antibodies were conjugated in-house using the MaxPar X8 antibody labeling Kit (Fluidigm Sciences) according to manufactures instructions. For all non-cadmium metals or with the Maxpar MCP9 for cadmium metals, and respectively stored in Antibody Stabilization Buffer (Candor Bioscience GmbH) or HRP-Protector™ peroxidase stabilizer (Boca Scientific) was used. Samples were incubated with 1 μM Cell-ID intercalator-103Rh to identify dead cells, followed by FcR blockage with mouse serum (2%) and FcR blocking anti-mouse CD16/32 antibodies (clone 2.4G2, BD Biosciences). Next, the metal-conjugated antibody mix was added, and cells were incubated overnight up to 48 h with 125 nM Cell-ID Intercalator-Ir in MaxPar Fix and Perm. Prior to acquisition on a Helios mass cytometer (Fluidigm, San Francisco, CA, USA), samples were centrifuged and resuspended in MilliQ and measured directly. Data were normalized using EQ Four Element Calibration Beads with the reference EQ passport P13H2302. Data analysis was performed by pre-gating live singlet CD45^+^ cells using FlowJo software (Tree Star), followed by non-supervised clustering based using the hierarchical t-SNE (HSNE) function of Cytosplore with 5 levels. Downstream analysis was performed using Cytofast (46, 47).

### Single-Cell RNA Sequencing and data analysis with the Seurat Package

To purify mouse T cells from blood, the Pan T Cell Isolation Kit I for mouse (Miltenyi Biotec) was used two consecutive times to reach >98% T cell purity after red blood cell lysis. A consecutive step to remove debris from the blood was performed by using the Debris Removal Solution (Miltenyi Biotec) according to the manufacturer protocol. Next, cells were subjected to single-cell-RNA-sequencing. Droplet-based 3’ end massively parallel single-cell RNA sequencing (scRNAseq) was performed by encapsulating sorted live T cells into droplets and libraries were prepared using Chromium Single Cell 3’ Reagent Kits v1 according to manufacturer’s protocol (10x Genomics). The generated scRNAseq libraries were sequenced using a NextSeq500 instrument by GenomeScan (Leiden, The Netherlands) with a sequencing depth of at least 50,000 reads per cell.

Downstream analysis was performed using the Seurat R package (67). Briefly, for each sample, mitochondrial, ribosomal and hemoglobin genes were excluded. Further, cells expressing less than 200 genes, and genes that were expressed in less than 3 cells were excluded. Next, all samples were pooled together into one dataset, and outlier cells expressing more than 2900 genes were excluded, which resulted in a dataset of 5260 cells. Next, the dataset was log1p normalized with a scaling factor of 10,000. Next, the set of 1709 highly variable genes were selected for further analysis. Dataset was preprocessed using principal component analysis. Using the top 15 principle components, the dataset was clustered using Louvain (graph-based community detection) and visualized using tSNE (68). Differentially expressed genes were identified between different cell groups, using Wilcoxon rank sum test with Bonferroni multiple test correction (adjusted P-value < 0.05). Within the CD4^+^ and CD8^+^ T cell populations, cells expressing the *Id2, Klrk1* and *Klrc1* genes were compared. DE genes were obtained between positive and negative groups of cells expressing these genes (expression > 1 was considered positive, otherwise negative).

### Bulk RNA Sequencing and data analysis with Qlucore

Splenic CD43^1B11+^ and CD43^1B11-^ CD8^+^ T cells from PDOX treated mice were FACS sorted using the BD FACSAria. RNA was isolated and the libraries were rRNA-depleted and stranded. The libraries were prepped using the Kapa Hyper Prep kit and adapters from IDT containing an 8bp UMI sequence were used. Sequencing was performed on the Illumina platform Novaseq6000 by Genomescan. To align the reads STAR 2.7.3a was used and to remove UMI detected sequences Umitools version 0.5.5 was used. HTseq-count was used to quantify the reads to gene count. GRCm38 was used as reference genome and for gene annotation Ensemble mouse gene version 99 was used. Differential-expression analysis was performed in Qlucore Omics Explorer (version 3.7), using the trimmed mean of log expression ratios method (TMM). Differentially expressed genes were selected (q<0.05) and used for pathway analysis with IPA.

### Immunofluorescence

For immunofluorescence images 4 μm FFPE tissue sections were deparaffinized, endogenous peroxidase was blocked with hydrogen peroxide and heat induced epitope retrieval was performed in with citrate (10 mM, pH 6.0). SuperBlock (ThermoFisher Scientific) was used to block nonspecific binding sites. Images were acquired with the Vectra V.3.0.5 microscope (PerkinElmer) at 20× magnification after staining with fluorescently-labeled anti-CD8 and DAPI as nuclear counterstain. CD8^+^ T cells in the TME were automatically phenotyped and counted with inForm V.2.4 image analysis software (PerkinElmer), after manual training and validation of procedures.

### Gene Expression and protein interaction profiling

Gene expression profiling interactive analysis (69) was used to generate the TCGA-based survival and correlation data. The search tool for retrieval of interacting genes (STRING) was applied to predict functional interactions of proteins (70).

### Statistics

Survival of the differentially treated animals was compared by the Kaplan-Meier method and log-rank (Mantel-cox) test. Statistical analysis of in vivo tumor growth was performed with Kruskal-Wallis. Other statistical comparisons were performed using an unpaired two-tailed Student’s t test (for 2 groups) or ANOVA (>2 group comparisons). Differences were considered statistically significant at *p* < 0.05.

## References

1. Sharma P, Siddiqui BA, Anandhan S, Yadav SS, Subudhi SK, Gao J, et al. The Next Decade of Immune Checkpoint Therapy. Cancer Discov. 2021;11(4):838–57.

2. van der Leun AM, Thommen DS, Schumacher TN. CD8(+) T cell states in human cancer: insights from single-cell analysis. Nat Rev Cancer. 2020;20(4):218–32.

3. Hiam-Galvez KJ, Allen BM, Spitzer MH. Systemic immunity in cancer. Nat Rev Cancer. 2021;21(6):345–59.

4. Galon J, Costes A, Sanchez-Cabo F, Kirilovsky A, Mlecnik B, Lagorce-Pages C, et al. Type, density, and location of immune cells within human colorectal tumors predict clinical outcome. Science. 2006;313(5795):1960–4.

5. Pardoll DM. The blockade of immune checkpoints in cancer immunotherapy. Nat Rev Cancer. 2012;12(4):252–64.

6. Arens R. Rational design of vaccines: learning from immune evasion mechanisms of persistent viruses and tumors. Adv Immunol. 2012;114:217–43.

7. Robert C, Schachter J, Long GV, Arance A, Grob JJ, Mortier L, et al. Pembrolizumab versus Ipilimumab in Advanced Melanoma. N Engl J Med. 2015;372(26):2521–32.

8. Robert C, Ribas A, Schachter J, Arance A, Grob JJ, Mortier L, et al. Pembrolizumab versus ipilimumab in advanced melanoma (KEYNOTE-006): post-hoc 5-year results from an open-label, multicentre, randomised, controlled, phase 3 study. Lancet Oncol. 2019;20(9):1239–51.

9. Hargadon KM, Johnson CE, Williams CJ. Immune checkpoint blockade therapy for cancer: An overview of FDA-approved immune checkpoint inhibitors. Int Immunopharmacol. 2018;62:29–39.

10. Haslam A, Prasad V. Estimation of the Percentage of US Patients With Cancer Who Are Eligible for and Respond to Checkpoint Inhibitor Immunotherapy Drugs. JAMA Netw Open. 2019;2(5):e192535.

11. Benedict CA, Ware CF. Virus targeting of the tumor necrosis factor superfamily. Virology. 2001;289(1):1–5.

12. Shum B, Larkin J, Turajlic S. Predictive biomarkers for response to immune checkpoint inhibition. Semin Cancer Biol. 2021.

13. Darvin P, Toor SM, Sasidharan Nair V, Elkord E. Immune checkpoint inhibitors: recent progress and potential biomarkers. Exp Mol Med. 2018;50(12):1–11.

14. Nixon AB, Schalper KA, Jacobs I, Potluri S, Wang IM, Fleener C. Peripheral immune-based biomarkers in cancer immunotherapy: can we realize their predictive potential? Journal for ImmunoTherapy of Cancer. 2019;7(1):325.

15. Schaer DA, Hirschhorn-Cymerman D, Wolchok JD. Targeting tumor-necrosis factor receptor pathways for tumor immunotherapy. J Immunother Cancer. 2014;2:7.

16. Marin-Acevedo JA, Dholaria B, Soyano AE, Knutson KL, Chumsri S, Lou Y. Next generation of immune checkpoint therapy in cancer: new developments and challenges. J Hematol Oncol. 2018;11(1):39.

17. Gubin MM, Zhang X, Schuster H, Caron E, Ward JP, Noguchi T, et al. Checkpoint blockade cancer immunotherapy targets tumour-specific mutant antigens. Nature. 2014;515(7528):577–81.

18. Newell EW, Sigal N, Nair N, Kidd BA, Greenberg HB, Davis MM. Combinatorial tetramer staining and mass cytometry analysis facilitate T-cell epitope mapping and characterization. Nat Biotechnol. 2013;31(7):623–9.

19. Hartmann FJ, Babdor J, Gherardini PF, Amir ED, Jones K, Sahaf B, et al. Comprehensive Immune Monitoring of Clinical Trials to Advance Human Immunotherapy. Cell Rep. 2019;28(3):819–31 e4.

20. Guruprasad P, Lee YG, Kim KH, Ruella M. The current landscape of single-cell transcriptomics for cancer immunotherapy. J Exp Med. 2021;218(1).

21. Gubin MM, Esaulova E, Ward JP, Malkova ON, Runci D, Wong P, et al. High-Dimensional Analysis Delineates Myeloid and Lymphoid Compartment Remodeling during Successful Immune-Checkpoint Cancer Therapy. Cell. 2018;175(4):1014-30.e19.

22. Li H, van der Leun AM, Yofe I, Lubling Y, Gelbard-Solodkin D, van Akkooi ACJ, et al. Dysfunctional CD8 T Cells Form a Proliferative, Dynamically Regulated Compartment within Human Melanoma. Cell. 2019;176(4):775-89.e18.

23. Sade-Feldman M, Yizhak K, Bjorgaard SL, Ray JP, de Boer CG, Jenkins RW, et al. Defining T Cell States Associated with Response to Checkpoint Immunotherapy in Melanoma. Cell. 2018;175(4):998-1013.e20.

24. Felix J, Lambert J, Roelens M, Maubec E, Guermouche H, Pages C, et al. Ipilimumab reshapes T cell memory subsets in melanoma patients with clinical response. Oncoimmunology. 2016;5(7):1136045.

25. Wistuba-Hamprecht K, Martens A, Heubach F, Romano E, Geukes Foppen M, Yuan J, et al. Peripheral CD8 effector-memory type 1 T-cells correlate with outcome in ipilimumab-treated stage IV melanoma patients. Eur J Cancer. 2017;73:61–70.

26. Manjarrez-Orduno N, Menard LC, Kansal S, Fischer P, Kakrecha B, Jiang C, et al. Circulating T Cell Subpopulations Correlate With Immune Responses at the Tumor Site and Clinical Response to PD1 Inhibition in Non-Small Cell Lung Cancer. Front Immunol. 2018;9:1613.

27. Takeuchi Y, Tanemura A, Tada Y, Katayama I, Kumanogoh A, Nishikawa H. Clinical response to PD-1 blockade correlates with a sub-fraction of peripheral central memory CD4+ T cells in patients with malignant melanoma. Int Immunol. 2018;30(1):13–22.

28. Martens A, Wistuba-Hamprecht K, Yuan J, Postow MA, Wong P, Capone M, et al. Increases in Absolute Lymphocytes and Circulating CD4+ and CD8+ T Cells Are Associated with Positive Clinical Outcome of Melanoma Patients Treated with Ipilimumab. Clin Cancer Res. 2016;22(19):4848–58.

29. Wu TD, Madireddi S, de Almeida PE, Banchereau R, Chen YJ, Chitre AS, et al. Peripheral T cell expansion predicts tumour infiltration and clinical response. Nature. 2020;579(7798):274–8.

30. Jacquelot N, Roberti MP, Enot DP, Rusakiewicz S, Ternès N, Jegou S, et al. Predictors of responses to immune checkpoint blockade in advanced melanoma. Nat Commun. 2017;8(1):592.

31. Wang W, Yu D, Sarnaik AA, Yu B, Hall M, Morelli D, et al. Biomarkers on melanoma patient T cells associated with ipilimumab treatment. J Transl Med. 2012;10:146.

32. Yamauchi T, Hoki T, Oba T, Jain V, Chen H, Attwood K, et al. T-cell CX3CR1 expression as a dynamic blood-based biomarker of response to immune checkpoint inhibitors. Nat Commun. 2021;12(1):1402.

33. Fairfax BP, Taylor CA, Watson RA, Nassiri I, Danielli S, Fang H, et al. Peripheral CD8(+) T cell characteristics associated with durable responses to immune checkpoint blockade in patients with metastatic melanoma. Nat Med. 2020;26(2):193–9.

34. Subrahmanyam PB, Dong Z, Gusenleitner D, Giobbie-Hurder A, Severgnini M, Zhou J, et al. Distinct predictive biomarker candidates for response to anti-CTLA-4 and anti-PD-1 immunotherapy in melanoma patients. J Immunother Cancer. 2018;6(1):18.

35. Krieg C, Nowicka M, Guglietta S, Schindler S, Hartmann FJ, Weber LM, et al. High-dimensional single-cell analysis predicts response to anti-PD-1 immunotherapy. Nat Med. 2018;24(2):144–53.

36. Martens A, Wistuba-Hamprecht K, Geukes Foppen M, Yuan J, Postow MA, Wong P, et al. Baseline Peripheral Blood Biomarkers Associated with Clinical Outcome of Advanced Melanoma Patients Treated with Ipilimumab. Clin Cancer Res. 2016;22(12):2908–18.

37. Spitzer MH, Nolan GP. Mass Cytometry: Single Cells, Many Features. Cell. 2016;165(4):780–91.

38. Corbett TH, Griswold DP, Jr., Roberts BJ, Peckham JC, Schabel FM, Jr. Tumor induction relationships in development of transplantable cancers of the colon in mice for chemotherapy assays, with a note on carcinogen structure. Cancer Res. 1975;35(9):2434–9.

39. Sagiv-Barfi I, Czerwinski DK, Levy S, Alam IS, Mayer AT, Gambhir SS, et al. Eradication of spontaneous malignancy by local immunotherapy. Science translational medicine. 2018;10(426):eaan4488.

40. Bald T, Quast T, Landsberg J, Rogava M, Glodde N, Lopez-Ramos D, et al. Ultraviolet-radiation-induced inflammation promotes angiotropism and metastasis in melanoma. Nature. 2014;507(7490):109–13.

41. Cannarile MA, Lind NA, Rivera R, Sheridan AD, Camfield KA, Wu BB, et al. Transcriptional regulator Id2 mediates CD8+ T cell immunity. Nat Immunol. 2006;7(12):1317–25.

42. Omilusik KD, Nadjsombati MS, Shaw LA, Yu B, Milner JJ, Goldrath AW. Sustained Id2 regulation of E proteins is required for terminal differentiation of effector CD8(+) T cells. J Exp Med. 2018;215(3):773–83.

43. Correia MP, Costa AV, Uhrberg M, Cardoso EM, Arosa FA. IL-15 induces CD8+ T cells to acquire functional NK receptors capable of modulating cytotoxicity and cytokine secretion. Immunobiology. 2011;216(5):604–12.

44. Harrington LE, Galvan M, Baum LG, Altman JD, Ahmed R. Differentiating between memory and effector CD8 T cells by altered expression of cell surface O-glycans. J Exp Med. 2000;191(7):1241–6.

45. van Unen V, Hollt T, Pezzotti N, Li N, Reinders MJT, Eisemann E, et al. Visual analysis of mass cytometry data by hierarchical stochastic neighbour embedding reveals rare cell types. Nat Commun. 2017;8(1):1740.

46. Beyrend G, Stam K, Hollt T, Ossendorp F, Arens R. Cytofast: A workflow for visual and quantitative analysis of flow and mass cytometry data to discover immune signatures and correlations. Comput Struct Biotechnol J. 2018;16:435–42.

47. Beyrend G, Stam K, Ossendorp F, Arens R. Visualization and Quantification of High-Dimensional Cytometry Data using Cytofast and the Upstream Clustering Methods FlowSOM and Cytosplore. J Vis Exp. 2019(154).

48. Prajapati K, Perez C, Rojas LBP, Burke B, Guevara-Patino JA. Functions of NKG2D in CD8(+) T cells: an opportunity for immunotherapy. Cell Mol Immunol. 2018;15(5):470–9.

49. Matsumoto M, Atarashi K, Umemoto E, Furukawa Y, Shigeta A, Miyasaka M, et al. CD43 functions as a ligand for E-Selectin on activated T cells. J Immunol. 2005;175(12):8042–50.

50. Piali L, Weber C, LaRosa G, Mackay CR, Springer TA, Clark-Lewis I, et al. The chemokine receptor CXCR3 mediates rapid and shear-resistant adhesion-induction of effector T lymphocytes by the chemokines IP10 and Mig. Eur J Immunol. 1998;28(3):961–72.

51. Wang R, Gao C, Raymond M, Dito G, Kabbabe D, Shao X, et al. An Integrative Approach to Inform Optimal Administration of OX40 Agonist Antibodies in Patients with Advanced Solid Tumors. 2019.

52. Brahmer JR, Tykodi SS, Chow LQ, Hwu WJ, Topalian SL, Hwu P, et al. Safety and activity of anti-PD-L1 antibody in patients with advanced cancer. N Engl J Med. 2012;366(26):2455–65.

53. Messenheimer DJ, Jensen SM, Afentoulis ME, Wegmann KW, Feng Z, Friedman DJ, et al. Timing of PD-1 Blockade Is Critical to Effective Combination Immunotherapy with Anti-OX40. Clin Cancer Res. 2017.

54. Ma Y, Li J, Wang H, Chiu Y, Kingsley CV, Fry D, et al. Combination of PD1 Inhibitor and OX40 Agonist Induces Tumor Rejection and Immune Memory in Mouse Models of Pancreatic Cancer. Gastroenterology. 2020.

55. Bansal-Pakala P, Halteman BS, Cheng MH, Croft M. Costimulation of CD8 T cell responses by OX40. J Immunol. 2004;172(8):4821–5.

56. Rogers PR, Song J, Gramaglia I, Killeen N, Croft M. OX40 promotes Bcl-xL and Bcl-2 expression and is essential for long-term survival of CD4 T cells. Immunity. 2001;15(3):445–55.

57. Ahrends T, Spanjaard A, Pilzecker B, Babala N, Bovens A, Xiao Y, et al. CD4(+) T Cell Help Confers a Cytotoxic T Cell Effector Program Including Coinhibitory Receptor Downregulation and Increased Tissue Invasiveness. Immunity. 2017;47(5):848-61.e5.

58. Herndler-Brandstetter D, Ishigame H, Shinnakasu R, Plajer V, Stecher C, Zhao J, et al. KLRG1(+) Effector CD8(+) T Cells Lose KLRG1, Differentiate into All Memory T Cell Lineages, and Convey Enhanced Protective Immunity. Immunity. 2018;48(4):716-29.e8.

59. van Duikeren S, Fransen MF, Redeker A, Wieles B, Platenburg G, Krebber WJ, et al. Vaccine-induced effector-memory CD8+ T cell responses predict therapeutic efficacy against tumors. J Immunol. 2012;189(7):3397–403.

60. Mathewson ND, Ashenberg O, Tirosh I, Gritsch S, Perez EM, Marx S, et al. Inhibitory CD161 receptor identified in glioma-infiltrating T cells by single-cell analysis. Cell. 2021;184(5):1281-98.e26.

61. Kagamu H, Kitano S, Yamaguchi O, Yoshimura K, Horimoto K, Kitazawa M, et al. CD4(+) T-cell Immunity in the Peripheral Blood Correlates with Response to Anti-PD-1 Therapy. Cancer Immunol Res. 2020;8(3):334–44.

62. Borst L, van der Burg SH, van Hall T. The NKG2A-HLA-E Axis as a Novel Checkpoint in the Tumor Microenvironment. Clin Cancer Res. 2020;26(21):5549–56.

63. Fuertes MB, Domaica CI, Zwirner NW. Leveraging NKG2D Ligands in Immuno-Oncology. Front Immunol. 2021;12:713158.

64. André P, Denis C, Soulas C, Bourbon-Caillet C, Lopez J, Arnoux T, et al. Anti-NKG2A mAb Is a Checkpoint Inhibitor that Promotes Anti-tumor Immunity by Unleashing Both T and NK Cells. Cell. 2018;175(7):1731-43.e13.

65. Beyrend G, van der Gracht E, Yilmaz A, van Duikeren S, Camps M, Hollt T, et al. PD-L1 blockade engages tumor-infiltrating lymphocytes to co-express targetable activating and inhibitory receptors. J Immunother Cancer. 2019;7(1):217.

66. Friedman CF, Postow MA. Emerging Tissue and Blood-Based Biomarkers that may Predict Response to Immune Checkpoint Inhibition. Curr Oncol Rep. 2016;18(4):21.

67. Butler A, Hoffman P, Smibert P, Papalexi E, Satija R. Integrating single-cell transcriptomic data across different conditions, technologies, and species. Nat Biotechnol. 2018;36(5):411–20.

68. van der Maaten L, Hinton, G. Visualizing Data using t-SNE. Journal of Machine Learning Research. 2008.

69. Tang Z, Li C, Kang B, Gao G, Li C, Zhang Z. GEPIA: a web server for cancer and normal gene expression profiling and interactive analyses. Nucleic Acids Res. 2017;45(W1):W98–w102.

70. Szklarczyk D, Franceschini A, Wyder S, Forslund K, Heller D, Huerta-Cepas J, et al. STRING v10: protein-protein interaction networks, integrated over the tree of life. Nucleic Acids Res. 2015;43(Database issue):D447–52.

