## Supplemental Figures for "OX40 agonism enhances efficacy of PD-L1 checkpoint blockade by shifting the cytotoxic T cell differentiation spectrum"

#### Supplementary Figures

**Figure S1.**

(A) Schematic of the strategy. WT mice were subcutaneously challenged with MC-38 syngenic tumors and were left untreated or treated with anti-OX40, CpG or anti-OX40/CpG.

(B) Tumor growth (mean ± SEM) of untreated and treated mice. One of 2 independent experiments is shown.

(C) Volcano plots showing significant gene expression related to Klrk1, and Klrc1 expression in $CD4^{+}$ and $CD8^{+}$ T cells. The log2 fold change (FC) in gene expression on the x-axis and unadjusted P values on the y-axis are illustrated. Black dots represent genes with adjusted P-value > 0.05, red dots represents genes with adjusted P-value < 0.05 and absolute average log2 Fold-Change < 1, green dots with gene name represent genes with adjusted P-value < 0.05 and absolute average log2 Fold-Change > 1. Bar graphs indicate the percentage of the cell origin according to their treatment.

**Figure S2.**

(A) Percentage of $CD43^{S7+}$ and $CD43^{1B11+}$ cells within $CD8^{+}$ T cells in blood of untreated and PDOX treated mice. Data are represented as mean ± SEM. *p<0.05, **p<0.01, ***p<0.001, by unpaired student's t test. Each dot represents an individual mouse.

(B) Percentage of $CD43^{1B11+}$ cells within the total $CD8^{+}$ T cell population in blood of untreated and ICT treated groups. Data are represented as mean ± SEM. *p<0.05, **p<0.01, ***p<0.001, by unpaired student's t test. Each dot represents an individual mouse.

(C) Representative histograms of marker expression in blood circulating $CD43^{1B11+}$ and $CD43^{1B11-}$ $CD8^{+}$ and $CD43^{1B11+}$ and $CD43^{1B11-}$ $CD4^{+}$ T cell populations of MC-38 challenged PDOX treated mice.

**Figure S3.**

Mass cytometry panels. Mouse and human panel used to analyze either the blood, spleen, lymph nodes and bone marrow from mice or blood of patients.

**Figure S4.**

Heatmaps of selected T cell clusters in blood, bone marrow, spleen and lymph node of untreated and ICT treated mice. The level of ArcSinh5 transformed marker expression is displayed by a rainbow scale. Bar graphs indicate the abundance and significant differences of the selected T cell clusters. Data are represented as mean ± SEM. *p<0.05, **p<0.01, ***p<0.001.

**Figure S5.**

Percentage CD43$^{1B11+}$ cells of CD8$^{+}$ T cells in blood and percentage CD43$^{1B11+}$ and NKG2A$^{+}$ CD8$^{+}$ T cells of live cells in the tumor-micro-environment of untreated and PDOX treated mice. *p<0.05, **p<0.01, ***p<0.001

**Figure S6.**

(A) The Cancer Genome Atlas (TCGA) survival plots for high vs low *KLRB1* expression for adrenocortical carcinoma (ACC) and sarcoma (SARC).

(B) TCGA survival and correlation analysis of *GZMB* expression for skin cutaneous melanoma (SKCM). Spearman correlation coefficient is indicated.

### Figure S1

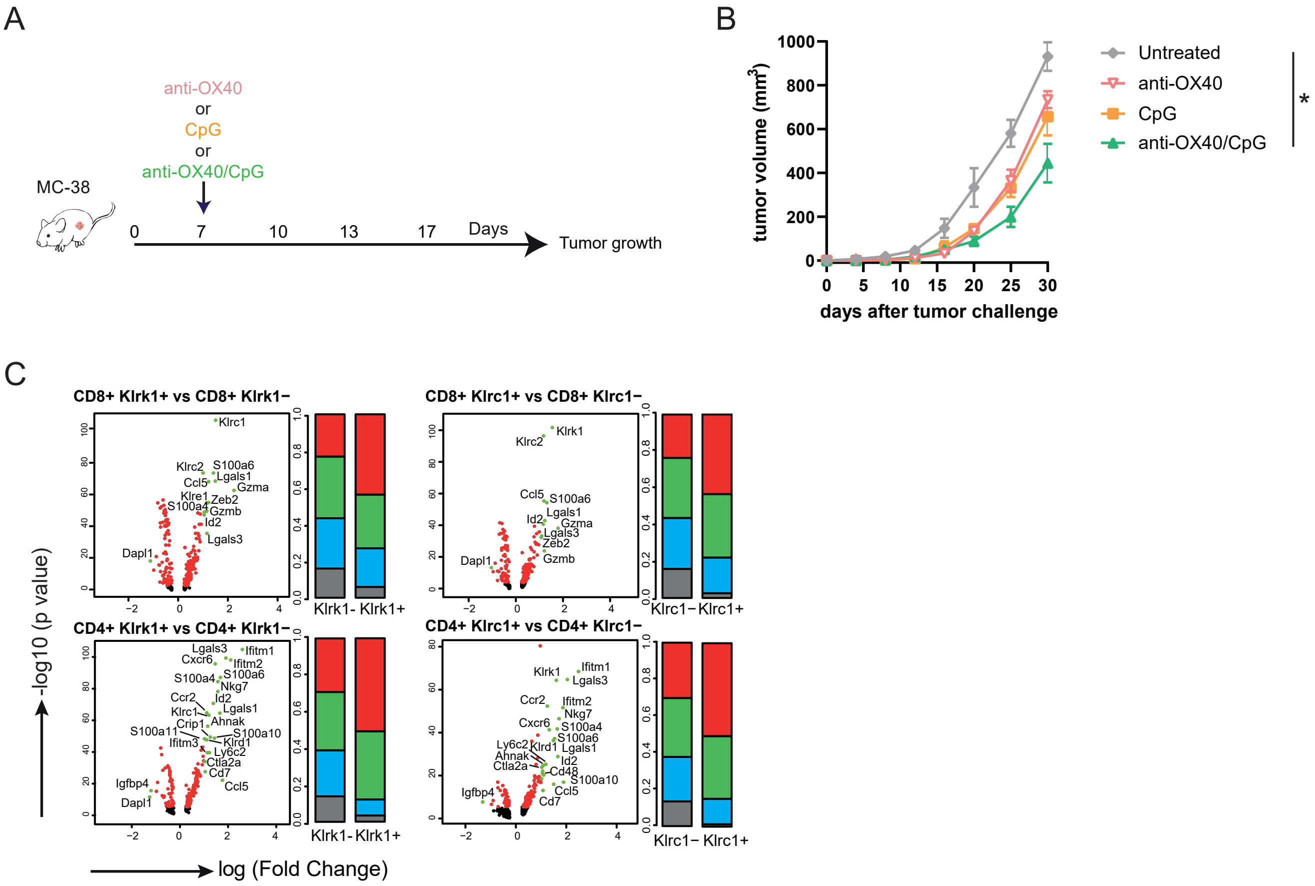

A
anti-OX40
or
CpG
or
anti-OX40/CpG
MC-38
0
7
10
13
17
Days
Tumor growth
B
tumor volume (mm³)
days after tumor challenge
Untreated
anti-OX40
CpG
anti-OX40/CpG
*
C
CD8+ Klrk1+ vs CD8+ Klrk1−
CD8+ Klrc1+ vs CD8+ Klrc1−
CD4+ Klrk1+ vs CD4+ Klrk1−
CD4+ Klrc1+ vs CD4+ Klrc1−
Klrk1- Klrk1+
Klrc1− Klrc1+
-log10 (p value)
log (Fold Change)

Figure S2

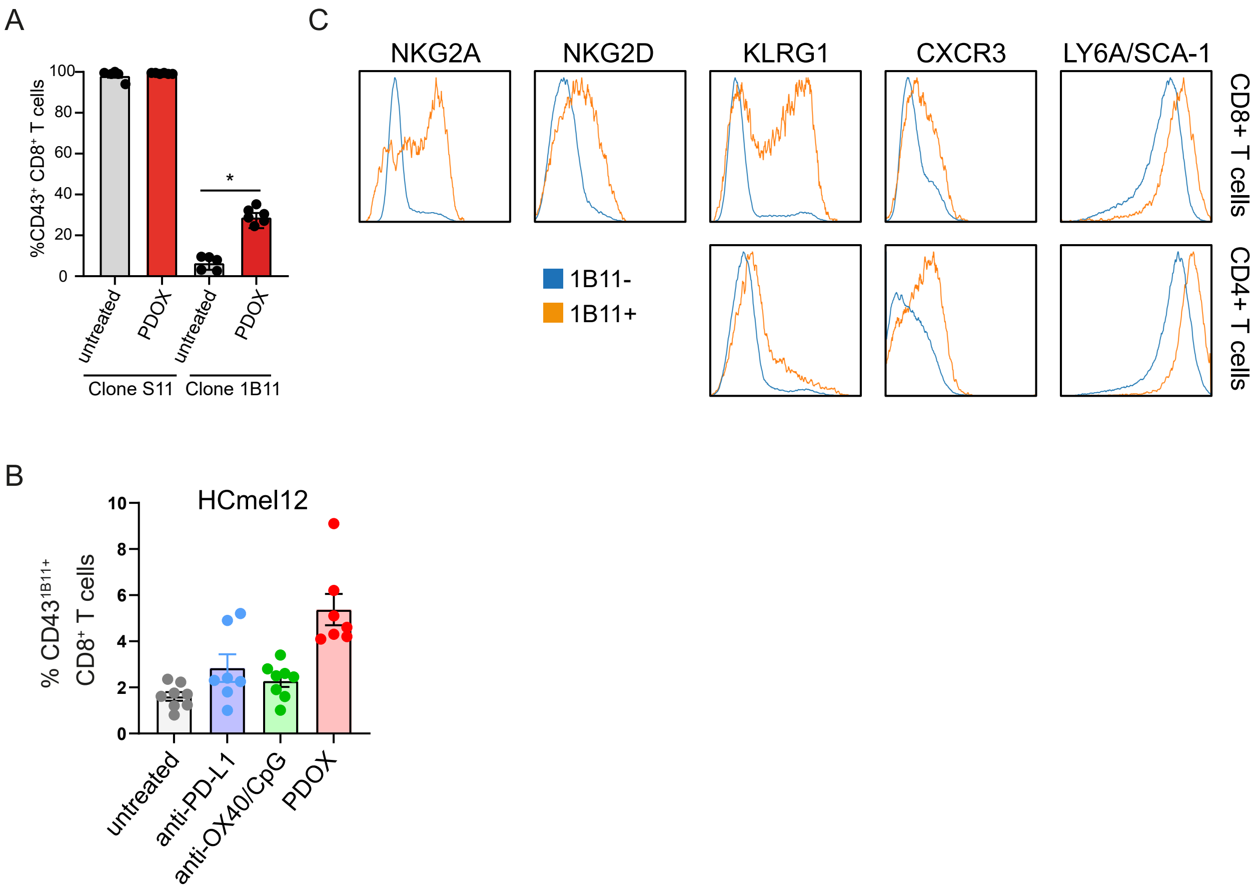

A
100
80
60
40
20
0
%CD43+ CD8+ T cells
*
untreated
PDOX
untreated
PDOX
Clone S11
Clone 1B11
B
HCmel12
10
8
6
4
2
0
% CD43 1B11+ CD8+ T cells
untreated
anti-PD-L1
anti-OX40/CpG
PDOX
C
NKG2A
NKG2D
KLRG1
CXCR3
LY6A/SCA-1
CD8+ T cells
CD4+ T cells
1B11-
1B11+

Figure S3

| Mouse panel | | |
|---|---|---|
| **Antigen** | **Clone** | **Company** |
| CD3e | 145-2C11 | eBiosciences |
| CD4 | RMA45 | Fluidigm |
| CD8A | 53-6.7 | Fluidigm |
| CD9 | eBioKMC8 | eBiosciences |
| CD11B | M1/70 | Fluidigm |
| CD11C | N418 | eBiosciences |
| CD14 | Sa14-2 | BioLegend |
| CD19 | 6D5 | Fluidigm |
| CD25 | 3C7 | Fluidigm |
| CD27 | LG.3A10 | eBiosciences |
| CD28 | 37.51 | Fluidigm |
| CD38 | 90 | eBiosciences |
| CD39 | 24DMS1 | eBiosciences |
| CD43 | 1B11 | BioLegend |
| CD44 | IM7 | eBiosciences |
| CD45 | 30-F11 | Fluidigm |
| ICAM-1 | YN1/1.7.4 | BioLegend |
| L-selectin | MEL-14 | BioLegend |
| CD69 | H1.2F3 | Fluidigm |
| B7-2 | GL-1 | eBiosciences |
| C-FMS | AFS98 | Fluidigm |
| CD122 | TM-b1 | BioLegend |
| CD127 | A7R34 | Fluidigm |
| NK1.1 | PK136 | eBiosciences |
| CXCR3 | CXCR3-173 | eBiosciences |
| CXCR5 | L138D7 | BioLegend |
| LAG-3 | eBioC9B7W | eBiosciences |
| PDL1 | 10F.9G2 | BioLegend |
| ICOS | 7E.17G9 | BioLegend |
| PD1 | 29F.1A12 | eBiosciences |
| F4/80 | BM8 | eBiosciences |
| KLRG1 | 2F1 | eBiosciences |
| Ly6C | HK1.4 | eBiosciences |
| Ly6G | 1A8 | Fluidigm |
| MHC II | M5/114.15.2 | eBiosciences |
| NKG2A | 20d5 | eBiosciences |
| TCRab | H57-597 | BioLegend |
| TCRgd | eBioGL3 | eBiosciences |

| Human panel | | |
|---|---|---|
| **Antigen** | **Clone** | **Company** |
| CD38 | HIT2 | BioLegend |
| CD45RO | UCHL1 | Fluidigm |
| CD45RA | HI100 | BioLegend |
| CD44 | IM7 | Fluidigm |
| CD45 | HI30 | Fluidigm |
| CD49b | P1E6-C5 | BioLegend |
| CD39 | A1 | BioLegend |
| HLA-DR | L243 | Fluidigm |
| CD29 | TS2/12 | BioLegend |
| CD11b (Mac-1) | ICRF44 | Fluidigm |
| TCRgd | 11F2 | Dianova |
| CD274 (PD-L1) | 29E.2A3 | Fluidigm |
| CD8a | RPA-T8 | Fluidigm |
| CD278/ICOS | C398.4A | Fluidigm |
| CD103 | Ber-ACT8 | BioLegend |
| CD49A | SR84 | BD Biosciences |
| CD27 | L128 | Fluidigm |
| CD127 (IL-7Ra) | A019D5 | Fluidigm |
| CD25 (IL-2R) | 2A3 | Fluidigm |
| CD3 | UCHT1 | Fluidigm |
| CD4 | RPA-T4 | Fluidigm |
| TIGIT | MBSA43 | eBioscience |
| CXCR5 | MAB190 | RnDsytems |
| CD62L | DREG-56 | BioLegend |
| CD69 | FN50 | BioLegend |
| CD86 | IT2.2 | Fluidigm |
| CD154 (CD40L) | 24-31 | ThermoFischer |
| CD134 (OX40) | ACT35 | BD Biosciences |
| CD161 | HP-3G10 | Fluidigm |
| CD335 (NKp46) | BAB281 | Fluidigm |
| KLRG1 | SA231A2 | BioLegend |
| CD223/LAG-3 | 11C3C65 | Fluidigm |
| CD20 | 2H7 | BioLegend |
| CD14 | Tük4 | ThermoFischer |
| CD56 | 5.1H11 | BioLegend |
| CD150 | A12 | BioLegend |
| CD244 | 2-69 | BioLegend |
| CD160 | BY55 | BioLegend |
| CCR6 | GE034E3 | BioLegend |
| CD122 | Tu27 | BioLegend |
| 4-1BB | 4B4-1 | BioLegend |
| CXCR6 | KE041E5 | BioLegend |
| NKG2A | 131411 | RnDsytems |
| PD-1 | EH12.2H7 | Fluidigm |

Figure S4

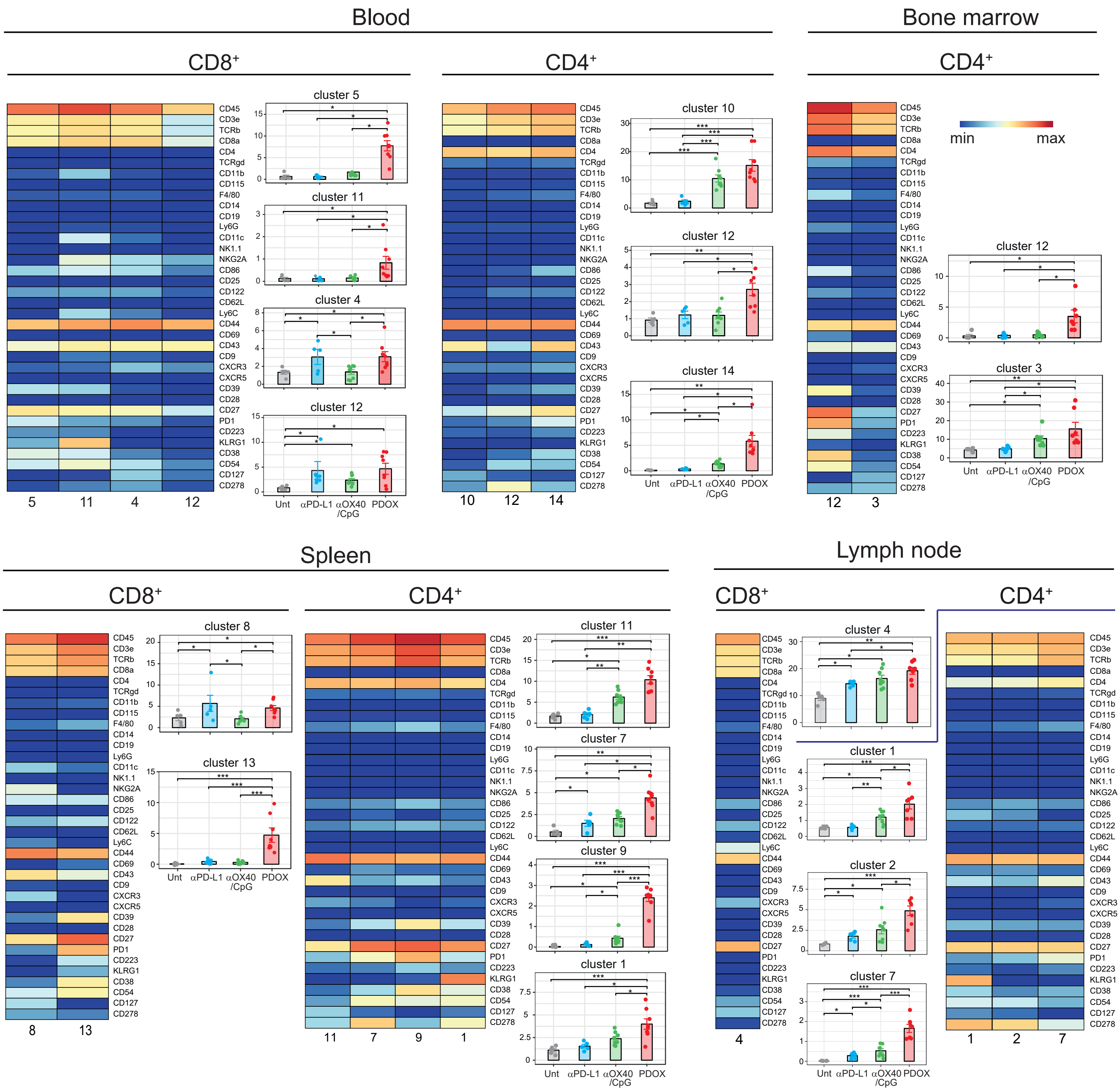

Blood
Bone marrow
CD8⁺
CD4⁺
CD4⁺
min
max
cluster 5
cluster 11
cluster 4
cluster 12
cluster 10
cluster 12
cluster 14
cluster 12
cluster 3
5
11
4
12
10
12
14
12
3
Spleen
Lymph node
CD8⁺
CD4⁺
CD8⁺
CD4⁺
cluster 8
cluster 13
cluster 11
cluster 7
cluster 9
cluster 1
cluster 4
cluster 1
cluster 2
cluster 7
8
13
11
7
9
1
4
1
2
7
CD45
CD3e
TCRb
CD8a
CD4
TCRgd
CD11b
CD115
F4/80
CD14
CD19
Ly6G
CD11c
NK1.1
NKG2A
CD86
CD25
CD122
CD62L
Ly6C
CD44
CD69
CD43
CD9
CXCR3
CXCR5
CD39
CD28
CD27
PD1
CD223
KLRG1
CD38
CD54
CD127
CD278
Unt
αPD-L1
αOX40 /CpG
PDOX

Figure S5

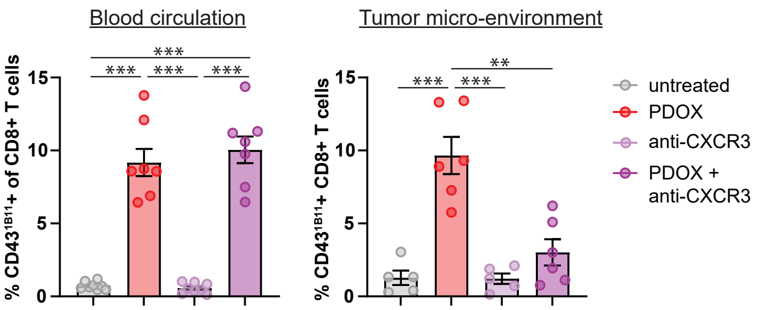
Blood circulation
Tumor micro-environment
% CD43[1B11]+ of CD8+ T cells
% CD43[1B11]+ CD8+ T cells
15
10
5
0
***
**
untreated
PDOX
anti-CXCR3
PDOX +
anti-CXCR3

Figure S6

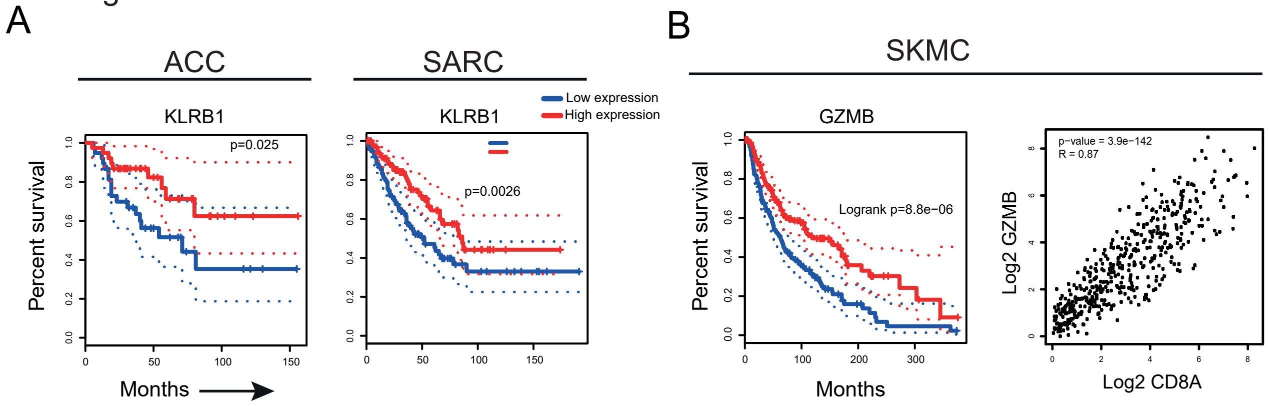
A
ACC
KLRB1
p=0.025
Percent survival
Months
SARC
KLRB1
Low expression
High expression
p=0.0026
B
SKMC
GZMB
Logrank p=8.8e−06
Percent survival
Months
p−value = 3.9e−142
R = 0.87
Log2 GZMB
Log2 CD8A
